# Two functional forms of the Meckel-Gruber syndrome protein TMEM67 generated by proteolytic cleavage by ADAMTS9 mediate Wnt signaling and ciliogenesis

**DOI:** 10.1101/2024.09.04.611229

**Authors:** Manu Ahmed, Sydney Fischer, Karyn L. Robert, Karen I. Lange, Michael W. Stuck, Sunayna Best, Colin A. Johnson, Gregory J. Pazour, Oliver E. Blacque, Sumeda Nandadasa

## Abstract

*TMEM67* mutations are the major cause of Meckel-Gruber syndrome. TMEM67 is involved in both ciliary transition zone assembly, and non-canonical Wnt signaling mediated by its extracellular domain. How TMEM67 performs these two separate functions is not known. We identify a novel cleavage motif in the extracellular domain of TMEM67 cleaved by the extracellular matrix metalloproteinase ADAMTS9. This cleavage regulates the abundance of two functional forms: A C-terminal portion which localizes to the ciliary transition zone regulating ciliogenesis, and a non-cleaved form which regulates Wnt signaling. By characterizing three *TMEM67* ciliopathy patient variants within the cleavage motif utilizing mammalian cell culture and *C. elegans,* we show the cleavage motif is essential for cilia structure and function, highlighting its clinical significance. We generated a novel non-cleavable TMEM67 mouse model which develop severe ciliopathies phenocopying *Tmem67*^-/-^ mice, but in contrast, undergo normal Wnt signaling, substantiating the existence of two functional forms of TMEM67.

## INTRODUCTION

Ciliopathies are a class of multi-organ developmental disorders caused by mutations affecting cilia formation or function. Frequently termed the ‘antennae of the cell, primary cilia are immotile, singular, microtubule-based signaling organelles that are present on almost all mammalian cell types [1–3]. The ciliary membrane is enriched with receptors that transduce signals in response to extracellular cues. These pattern the events of cellular differentiation, proliferation, and polarity, which together control tissue morphogenesis and organ formation during embryonic and postnatal development [4, 5]. Motile cilia on the other hand can be both singular (node cilia), or formed in multiciliated epithelia which regulate fluid flow dynamics in a variety of embryonic and postnatal tissue [6, 7]. All cilia possess a specialized diffusion barrier, termed the transition zone (TZ), found at the base of the cilium. The TZ acts as a “gatekeeper” to regulate molecular traffic between the cilium and the cytoplasm [8, 9]. The TZ is comprised of functional modules linked to ciliopathies, which are categorized based on genetic interactions and biochemical characterization, and include the Meckel-Gruber syndrome (MKS) and nephronophthisis (NPHP) modules. Despite genetic and phenotypic overlap, these complexes have distinct spatial locations and interaction networks within the TZ [10–12].

MKS, first described by Johann Friedrich Meckel in 1822, is a relatively rare autosomal recessive disorder, with a 100% mortality rate [13]. MKS is characterized by large polycystic kidneys, polydactyly, and occipital encephalocele and represents the most severe end of the ciliopathy disease spectrum in humans [13–15]. Pathogenic variants in the transmembrane protein TMEM67 are the most frequent cause of MKS, linked to 16-20% of all clinically diagnosed MKS cases. *TMEM67* mutations also result in the medullary cystic kidney disease NPHP and Joubert syndrome (JBTS), a severe neurodevelopmental disorder characterized by a pathognomonic cerebellar and brain stem malformation known as the molar tooth sign [16–20].

*TMEM67* variants also result in RHYNS syndrome, associated with retinitis pigmentosa, <u>hy</u>popituitarism, <u>n</u>ephronophthisis and <u>s</u>keletal dysplasia [21], as well as COACH syndrome, which is characterized by <u>c</u>erebellar vermis hypoplasia, <u>o</u>ligophrenia, <u>a</u>taxia, <u>c</u>oloboma, and <u>h</u>epatic defects [22–24]. The large number and the severity of these genetic disorders caused by *TMEM67* mutations underscores the central biological role played by TMEM67 in human development and health.

TMEM67 is a component of the MKS module of the ciliary TZ, which forms part of the transition zone “necklace”, composed of transmembrane and extracellular components of the TZ. These anchor the ciliary membrane to the Y linkers and the microtubule core, and form a functional diffusion barrier [8]. However, TMEM67 is also comprised of an extracellular cysteine-rich domain (CRD) at its very N-terminus which is homologous to the CRDs found in the Frizzled family receptors and the ROR family of receptor tyrosine kinases which bind the Wnt ligands, modulating both canonical and non-canonical Wnt signaling pathways [25]. Previous work has shown that TMEM67 CRD binds to Wnt5a and forms a complex with ROR2, a co-receptor for non-canonical Wnt signaling, resulting in ROR2 phosphorylation and the transmission of non-canonical Wnt signaling [26–29]. In the absence of TMEM67/ROR2-mediated non-canonical Wnt signaling, mouse brains and kidneys show highly elevated canonical Wnt signaling, and therefore TMEM67 function is crucial for regulating balanced canonical and non-canonical Wnt signaling during mouse development [26, 29]. How TMEM67 controls both TZ assembly and Wnt signaling and what factors govern each functionality of TMEM67 are unknown; and there has been a corresponding lack in our understanding of the molecular mechanism(s) underlying the ciliopathies caused by *TMEM67* loss of function in humans.

We previously identified the extracellular matrix metalloprotease ADAMTS9 (A Disintegrin and Metalloproteinase with Thrombospondin motifs, family member 9), as a novel ciliopathy locus, resulting in NPHP and JBTS [30]. ADAMTS9 is necessary for normal ciliogenesis and TZ assembly in humans and mice [31]. Catalytically active ADAMTS9 is highly concentrated at the base of the cilium in Rab11+ endocytic recycling vesicles, carrying cilia-bound cargo. This localization requires the cell surface receptors LRP-1/2, clathrin-mediated endocytosis and the recycling endosome to traffic ADAMTS9 into periciliary vesicles. ADAMTS9’s catalytic function is also crucial for ciliogenesis, as catalytically inactive ADAMTS9 does not rescue the loss of ciliogenesis of *ADAMTS9*-null RPE-1 cells [31]. However, the ADAMTS9 targets involved in ciliogenesis were not known.

To answer this question, we undertook an advanced proteomics screen utilizing the iTRAQ TAILS N-Terminomics technique [32], and identified novel ADAMTS9 substrates [33]. Here, we unveil two novel ADAMTS9 cleavage sites present in the N-terminal extracellular domain of TMEM67. We show that TMEM67 has two functional forms governed by this proteolytic cleavage, one regulating Wnt signaling and a second regulating ciliary TZ assembly. We present a novel non-cleavable *Tmem67* mouse model which is defective for ciliogenesis but in stark contrast to *Tmem67*-null mice, undergoes normal non-canonical and canonical Wnt signaling. Our work identifies a novel cellular mechanism in which the extracellular matrix degrading metalloproteinase ADAMTS9 modifies the function of TMEM67, from a non-canonical Wnt signaling co-receptor at the cell surface, to a TZ scaffold protein, here forth uncoupling the dual functionality of TMEM67 in Wnt signaling and cilia formation.

## METHODS

### Contact for reagent and resource sharing

Requests for resources, reagents, and further information should be directed to and will be fulfilled by the lead Contact, Dr. Sumeda Nandadasa

### Data availability

The data that support the findings of this study are available from the corresponding author upon reasonable request.

### Mice

All mouse experiments conducted in this study were carried out under the IACUC approved protocol (2021-0006). *Tmem67^+/-^* mice (C57BL/6NJ-*Tmem67^em1(IMPC)J^*/Mmjax) generated by the KOMP were purchased from the Jackson Laboratory (JAX, 051248) and were maintained in the C57BL/6J background. *Tmem67^1CLE/+^* founder mice were generated in the C57BL/6J background by Cyagen (Santa Clara, CA) animal modeling service upon contract, as outlined in **Sup.** Fig. 5A. In brief, targeted ES cell clones 2F7 and 1E10 were injected into C57BL/6 albino embryos, which were then implanted to CD-1 pseudopregnant females. Founder animals were identified using coat color and germline transmission was confirmed by breeding with C57BL/6J female mice and genotyping the resulting offspring. Two heterozygous male mice and two heterozygous female mice were generated from clones 1E10 and 2F7 respectively, and were maintained as independent founder lines. PCR amplification and Sanger sequencing were used to verify the Neomycin cassette removal and correct mutagenesis of the target residues in both lines. Founder mice were bred for two additional generations with C57BL/6J male and female mice (JAX, 000664) prior to breeding heterozygous mice to generate homozygous *Tmem67^1CLE/1CLE^* mice.

### Mammalian cell culture

Wild type hTERT-RPE-1 (ATCC, CRL-4000), and CRISPR-Cas9 knockout cell lines for *TMEM67* [28] and *ADAMTS9* [31] were cultured in DMEM/F12 (Gibco;11330-032;) with 10% FBS and 200 U/mL penicillin/streptomycin (Gibco; 15140-122) and maintained at 37°C with 5% CO2. RPE-1 growth medium was supplemented with 0.01mg/mL hygromycin B (Invitrogen; 10687010;). Mouse embryonic fibroblasts were harvested from E13.5 embryos from timed mating and cultured for one week in DMEM/F12 medium containing 10% FBS and 200 U/mL penicillin/streptomycin and passaged once prior to immortalization with Simian virus 40 large T antigen (SV40LT), as previously described [34].

### Plasmid DNA constructs and transfections

The TMEM67 cleavage site mutations (S1, S2, and S1+S2) and ciliopathy patient variants (F343V, K329T, and L349S) were generated using the Q5 Site-Directed Mutagenesis Kit (NEB; E0554S) utilizing the full-length, human TMEM67 plasmid construct in the pCDNA3.1Myc/His vector backbone, originally cloned by the Johnson lab [28]. TMEM67 Δ342 and N-331 constructs were cloned into the HindIII(5’) and XhoI(3’) restriction enzyme sites of the pSecTag2C vector (Invitrogen, V900-20) following overhang PCR amplification of the human Fl-TMEM67-Myc construct using Phusion High-Fidelity Taq DNA Polymerase (NEB, M0530). Both constructs were cloned in frame to the N-terminal Ig κ-chain leader sequence (vector signal peptide) and the 6x His and Myc tag at the C-terminus. The TMEM67 endogenous signal peptide sequence (M^1^-A^36^) was not PCR amplified in generating the N-331 construct to prevent duplication of signal peptides and coded only for TMEM67 Q^37^-K^331^ while the TMEM67 Δ342 construct coded for F^343^-F^993^ of human TMEM67. Ligated DNA was transformed into DH5α competent *E. coli* (NEB;C2987H) and grown on LB plates containing 100µg/mL carbenicillin. Plasmid DNA was extracted with the ZymoPURE II plasmid midiprep kit (Zymo Research; D4200). All constructs were sequence verified to be free of any undesired mutations by Sanger sequencing. Plasmid DNA transfections were carried out using the PEI MAX transfection reagent (Polysciences; 24765). DNA and PEI MAX was combined at a final ratio of 1:3 and triturated with 100µl Opti-MEM reduced serum medium (Gibco; 31985-070). 300ng of DNA was transfected per well into 8-chamber slides (Corning; 354118). Transfected cells were incubated at 37°C, 5% CO2 for 6 hours or overnight.

### Immunostaining and fluorescence microscopy of cultured cells

For immunostaining analysis, RPE-1 or MEF cells were cultured in 8-well chamber slides in DMEM/F12 medium containing 10% FBS followed by serum starvation for 24 hours to induce ciliogenesis. For visualizing primary cilia, samples were fixed in fresh 4% paraformaldehyde in 0.1% PBS-Tween (PBST) at room temperature for 20 minutes. For immunolabelling of the transition zone markers, cells were washed once in PBS and fixed in ice-cold methanol for 5 minutes at −20°C. The fixed cells were washed three times in PBST and blocked in 5% normal goat serum made in PBST for 1 hour at room temperature. Primary and secondary antibodies were diluted in 5% normal goat serum in PBST (see Key Resources Table for antibodies and dilutions). Samples were incubated with primary antibodies at 4°C overnight and secondary antibodies for 2 hours at room temperature followed by 3x10 minute washes with PBST and mounted with Prolong gold mounting media with DAPI (Invitrogen, P36931). Confocal images were acquired using a Leica SP8 Laser Scanning confocal microscope with a x63 1.47 NA oil immersion objective, utilizing the HyD hybrid detectors. The SP8 lightning feature was used for super-resolution images of the transition zone collected at a 1000x magnification.

### Fluorescence microscopy quantifications and statistical analysis

Measurements of cilia length/frequency and intensity of the ciliary transition zone were quantified using NIH Image J (FIJI). Cell counter and line trace functions were used for measuring percentage of ciliated cells and cilium lengths. Pixel intensity of the transition zone markers were quantified by measuring the mean pixel intensity of an ROI and performing background subtraction. In brief, a boxed region of interest (ROI) was drawn directly above the CEP170 staining and transferred to the corresponding transition zone marker channel (red/ 568 channel), and the arbitrary mean pixel intensity for the ROI was measured. The “restore selection” function was used to transfer the same ROI to all transition zones analyzed. GraphPad Prism 10 software (La Jolla, CA) was used to determine statistical significance by one-way ANOVA for cilia length and frequency, and two-way ANOVA for quantifying transition zone intensity.

### Transmission and Scanning Electron Microscopy

Mouse brains and kidney were dissected in and immediately fixed in 2.5% glutaraldehyde with 1.6% paraformaldehyde in 0.1 M sodium cacodylate buffer (pH 7.2). Samples were processed and analyzed at the University of Massachusetts Chan Medical School Electron Microscopy core facility according to standard procedures. Briefly, fixed samples were moved into fresh 2.5% glutaraldehyde, 1.6% paraformaldehyde in 0.1 M sodium cacodylate buffer and left overnight at 4°C. The samples were then rinsed twice in the same fixation buffer and post-fixed with 1% osmium tetroxide for 1h at room temperature. Samples were washed twice with DH2O for 5 minutes and then dehydrated through a graded ethanol series of 20% increments and two changes in 100% ethanol. Samples were then infiltrated with two changes of 100% propylene oxide first and then with a propylene oxide and SPI-Pon 812 50%-50% resin mixture overnight. Three changes of fresh 100% SPI-Pon 812 resin were done before the samples were polymerized at 68°C in plastic capsules. Samples were oriented and 70nm ultra-thin sections were collected using a Leica EM UC7 ultramicrotome equipped with a Diatome ultra 45 diamond knife. Sections were placed on copper support grids and contrasted with lead citrate and uranyl acetate. Sections were examined using a FEI Tecani 12 BT with 120 kV accelerating voltage, and images were captured using a Gatan TEM CCD camera. Samples processed for SEM and bulk-frozen-fracture SEM (kidneys), were fixed as above, then dehydrated through a graded series of ethanol and either directly critical point dried or quickly immersed in liquid nitrogen, placed on a liquid nitrogen cooled block and fractured. The tissue pieces thawed in 100% ethanol and critical point dried. The samples were then mounted on aluminum SEM stubs and metal coated (Au/Pb, 60/40) and imaged on a FEI Quanta 200 MKII FeSEM.

### Western blotting

Cells cultured in 6-well plates were lysed using 500μl Transmembrane lysis buffer (FIVEphoton Biochemicals; TmPER-50) for TMEM67 detection, or ice-cold Pierce RIPA lysis buffer (ThermoFisher; 8901) for phospho ROR2 detection. Lysis buffer was supplemented with Pierce protease inhibitor complex with EDTA (ThermoFisher; A32953) and Pierce phosphatase inhibitor (ThermoFisher; A32957). 6X Laemmli sample buffer containing 9% 2-mercaptoethanol, 4.8% glycerol, 6% SDS and 0.03% bromophenol blue was used for boiling cell lysates or conditioned medium samples for 10 minutes at 100°C prior to SDS-PAGE analysis. 6% acrylamide gels were used for ROR2 phosphorylation detection and 7.5% acrylamide gels were used on all other experiments. Proteins were transferred to Immobilon-FL transfer membranes (EMD Milipore; IPFL00010) and blocked with 5% nonfat dry milk in 0.1% PBS-Tween for 1hr. Experiments detecting phospho ROR2 used LI-COR Intercept blocking buffer (Li-COR; 927-620001) for blocking and primary antibody incubation. In all other experiments, primary and secondary antibodies were diluted in 5% nonfat dry milk in 0.1% PBS-Tween. Primary antibodies were diluted 1:1000 and incubated at 4°C overnight. LI-COR secondary antibodies (LI-COR Biosciences, Lincoln, NE) were diluted 1:10,000 and incubated for 2 hours at room temperature (see Key Resources Table for antibodies and dilutions). A LI-COR Odyssey M scanner was used to image fluorescent western blot membranes.

### qRT-PCR

RNA was harvested from RPE-1 cells or MEFs cultured in 6-well culture plates or using whole mouse kidneys. Briefly, mRNA was harvested by lysing cells with 300µl or 500μl for minced whole kidneys in Trizol reagent and snap freezing in liquid nitrogen. Thawed samples were sonicated for 30 seconds followed by chloroform extraction and isopropanol precipitation. RNA pellets were washed with 70% ethanol and dissolved in ultra-pure ddH2O. 2μg of mRNA from each sample were used in cDNA synthesis using the High Capacity cDNA Reverse Transcription Kit (ThermoFisher; 4368814). A BioRad CFX connect real time PCR machine was used in combination with Bullseye EvaGreen qPCR master mix (Midsci; BEQPCR-R) for determining ΔCT values. qRT-PCR primers used in this study are provided in Supplemental Table-1. *Gapdh, *β*-Actin* or *18s* ribosomal RNA expression were used for normalizing samples. Each sample was run in duplicate with at least 3 experimental replicates per group. Fold change was quantified by determining the 2^(-Δ1Δ1CT) values in relation to the average expression value of the control group. Unpaired Student’s two-tailed *t*-tests were used to assess statistical significance and GraphPad Prizm (version 10.3.0) was used to generate bar graphs.

### Tissue histopathology and immunostaining analysis

For Haematoxylin and Eosin (H&E) staining, dissected kidney, heart and liver samples were fixed in 4% PFA made in PBST at 4°C overnight, washed three times in PBST, and paraffin embedded using standard tissue processing procedure. 7μm paraffin sections were collected using a Leica RM2155 motorized microtome, deparaffined and stained with Haematoxylin and Eosin and mounted in Cystoseal XYL medium (Adwin Scientific; NC9527349). For Masson’s Trichrome staining of livers, paraffin sections were deparaffined, refixed in Bouin’s fixative (Electron Microscopy Sciences (EMS); 26367-01) for 1hr at 56°C, stained with Weigert’s iron Haematoxylin A & B solutions (EMS; 26367-02 and −03), Biebrich scarlet solution (EMS; 26367-04), and Aniline blue (EMS; 26367-06) solutions and treated with acetic acid prior to mounting in Cytoseal XYL mounting medium. For immunostaining brain sections, dissected mouse brains were fixed overnight at 4°C in 4% PFA made in PBST, washed three times in PBS, and consecutively dehydrated in 15% sucrose and 30% sucrose overnight at 4°C. Samples were cryo embedded in OCT (Tissue-Tek; 4583), and 7μm cryo sections were collected using a Leica CM 1950 cryostat at −25°C. Thawed cryo sections were washed in PBS, blocked for 1hr in 5% normal goat serum in PBST at room temperature, and stained with primary antibodies overnight at 4°C, diluted in 5% normal goat serum in PBST (see Key Resources Table for antibodies and dilutions). Sections were washed thrice in PBST and incubated with secondary antibodies diluted in PBST for 2hrs at room temperature. Sections were washed thrice in PBST and sealed overnight in Prolong gold antifade mounting medium with DAPI (Invitrogen, P36931). A Leica SP8 confocal microscope was used for imaging brain sections, and a TissueGnostics SL slide scanner was used for imaging kidney and heart H&E stained sections. Liver H&E and Masson’s trichrome stained sections were imaged using a Zeis Axioplan widefield microscope equipped with a Leica DMC6200 color camera.

### *Caenorhabditis elegans* strains and maintenance

*C. elegans* worm strains were maintained at 20°C on NGM agar plates seeded with OP50 *E. coli* using standard husbandry techniques [35]. A list of worm strains generated and/or used in this study can be found in **Sup. Table-2**.

### CRISPR/Cas9 genome engineering in *C. elegans*

The F249V and ΔCLE (L^238^-A, F^239^-A, T^248^-A, F^249^-A) CRISPR mutants were generated by injecting the Cas9 ribonucleoprotein complex [36] into *nphp-4(tm925)* worms. Edited progeny were identified using an *unc-58* co-CRISPR strategy [37], and *unc-58* was sequenced in all CRISPR strains to ensure it was wild-type [38]. Worms that contained the engineered edit were identified using a PCR based approach [28]. *C. elegans* primers used in this study can be found in **Sup. Table-3**. Single-stranded oligonucleotides (Sigma) were used as repair templates to engineer precise edits. CRISPR reagents were purchased from IDT: Alt-R Cas9 Nuclease V3 (IDT; 1081058), Alt-R tracrRNA (IDT; 1072533), and custom-generated, gene specific Alt-R crRNA. crRNA and repair template sequences are listed in **Sup. Table-4**. All CRISPR mutants were confirmed by Sanger sequencing and outcrossed twice before analysis.

### *C. elegans* quantitative assays to assess cilia structure/function

Dye filling assays were performed with DiO (Invitrogen; D275) diluted in M9 (22 mM KH2PO4, 42 mM Na2HPO4, 85.5 mM NaCl, 1 mM MgSO4). Synchronised populations of young adult hermaphrodites were incubated for 1hr with 10ng/μl DiO solution then allowed to recover on NGM plates for 30 minutes before being mounted on 4% agarose pads with 40mM tetramisole (Sigma; L9756). Worms were visualised with a 20x objective on a Leica DM5000B epifluorescence microscope. Dye filling was quantified by counting the number of phasmid neurons with dye uptake. Sample size was more than 120 worms from at least three independent replicates. Roaming assays were performed by placing a single hermaphrodite on a fully seeded NGM place for 20hrs at 20°C. The worm was removed from the plate and a 5mm^2^ grid was placed under it to count how many squares the worm entered. All values were normalised to *wild type* (sample size at least 40 worms from three independent replicates). Chemotaxis plates (9cm petri dishes with 10ml of chemotaxis agar: 2% agar, 5mM KPO4 pH 6, 1mM CaCl2, 1mM MgSO4) were prepared 16-24 hrs before the experiment. Two spots were marked on opposite sides of the plates 1.5cm from the edge, and 1μl of 1M sodium azide (Sigma; S2002) was added to each spot. 1μl of ethanol (Honeywell; 32294) or 1:200 benzaldehyde (Sigma; B1334) diluted in ethanol was then added to the spots. Young adult hermaphrodites were washed and added to the centre of the plate. Excess liquid was removed, and the worms were counted after 1hr. The chemotaxis index was calculated as (b−c)/n, where b is the number of worms within 1.5cm of the benzaldehyde spot, c is the number of worms within 1.5cm of the ethanol control, and n is the total number of worms on the plate. Three independent replicates were performed with at least 24 assays performed for each genotype.

### Statistical analysis for *C. elegans* data

Statistical analysis was performed using GraphPad Prism (version 10.1.2.). A Shapiro-Wilk test was used to determine if data were normally distributed. For parametric data, one-way ANOVA with a post hoc Tukey test was used to calculate p-values. For non-parametric values, a Kruskall- Wallis test followed by a Dunn’s post hoc test was used to calculate p-values. Graphs were made using Microsoft Excel.

## RESULTS

### Identification of TMEM67 as a novel substrate of the extracellular matrix metalloprotease ADAMTS9

To identify novel proteolytic substrates of ADMTS9 involved in ciliogenesis, we carried out a N-terminomics proteomics study utilizing the Terminal Amine Isotopic Labeling of Substrates (TAILS) technique [32], comparing *Wild type* (Wt) and *ADAMTS9*-null RPE-1 cells [33]. This highly specialized advanced proteomics approach, allowed us to label the neo N-termini generated by proteolytic cleavage of a protein utilizing isobaric tags and quantify N-terminal labeled peptide abundance by LC-MS/MS after their specific enrichment using a highly-branched polyglycerol polymer (HPG-ALD) (**Sup.** Fig.1). Amongst a handful of known extracellular matrix substrates of ADAMTS9, the study revealed only one novel substrate known to be a structural component of the cilium, which was TMEM67. Mapping of the TMEM67 neo-peptide identified two highly conserved cleavage sites in the extracellular domain of the ciliary transition zone protein TMEM67 occurring at K^331^-F^332^ and N^342^-F^343^, which we named cleavage site 1 and 2 respectively (**Fig. 1A**). The F^332^ -labeled, non-tryptic 11 amino acid peptide released by the two cleavage events was either completely lost or present at significantly low abundance in *ADAMTS9* KO medium (**Fig. 1B**). Mapping the identified peptide in the AlphaFold model of TMEM67 revealed the cleavage to occur at the end of a long linker region predicted to be in between the cysteine rich domain (CRD) and the β-sheet rich domain (BRD) (**Fig. 1A**). The two cleavage sites and the 11 amino acid sequence in between were also highly conserved across mammals (**Fig. 1C**). During the course of this study, the cryo-electron structure for TMEM67 was resolved at 3.3 Å resolution [39]. This revealed a homodimeric structure with 4 dimerization interfaces, in which the N-terminal extracellular domain of one protomer interacted with the BRD and the CRD of the second protomer, forming an extracellular arch [39]. Modeling the N-terminomics-identified cleavage sites in this cryo-EM structure revealed the cleavage to occur at the very N-terminus of the first homodimer interface, releasing a CRD-CRD dimer when cleaved (**Sup.** Fig. 2A-B). The cryo-EM structure also revealed that the cleavage sites were in two previously uncharacterized, highly accessible, long linker regions present in between the CRD and BRD of the TMEM67 extracellular domain, which we have named linker-1 (L1) and linker-2 (L2) respectively (**Sup.** Fig. 2C-D, **Fig.1D**). These unique characteristics gave us high confidence in further characterizing and validating the N-terminomics identified cleavage sites of TMEM67 utilizing N- and C-terminal-specific antibodies (**Fig. 1D**). Proteomics data predicted that a 34kDa N-terminal fragment would be shed by TMEM67 extracellular domain cleavage. Western blotting of the conditioned medium and the cell layers of Wt and *ADAMTS9*-null RPE-1 cells [31] utilizing the TMEM67 commercial antibody, validated this predication and revealed two closely migrating bands present only in the Wt conditioned medium and absent in the *ADAMTS9*-null conditioned medium, while the respective cell lysates showed the full-length TMEM67 (∼120kDa) to be abundant only in the *ADAMTS9*-null cells (**Fig. 1E**). Immunostaining with N- or C-terminal-specific TMEM67 antibodies [27], in Wt and *ADAMTS9*-null RPE-1 cells showed that only the TMEM67 C-terminus co-localized with the mature basal body marker CEP170, which marks the sub-distal appendages, in serum starved Wt RPE-1 cells. Neither the TMEM67 N-terminus nor the C-terminus co-localized with the mature basal body in *ADAMTS9* deficient cells (**Fig.1 F-G**). Since both TMEM67 and ADAMTS9 are required for ciliogenesis in RPE-1 cells [28, 31], these data suggest TMEM67 cleavage may be a prerequisite for ciliogenesis and only the TMEM67 C-terminal half, generated by cleavage, may be present at the ciliary TZ.

**Figure 1:**
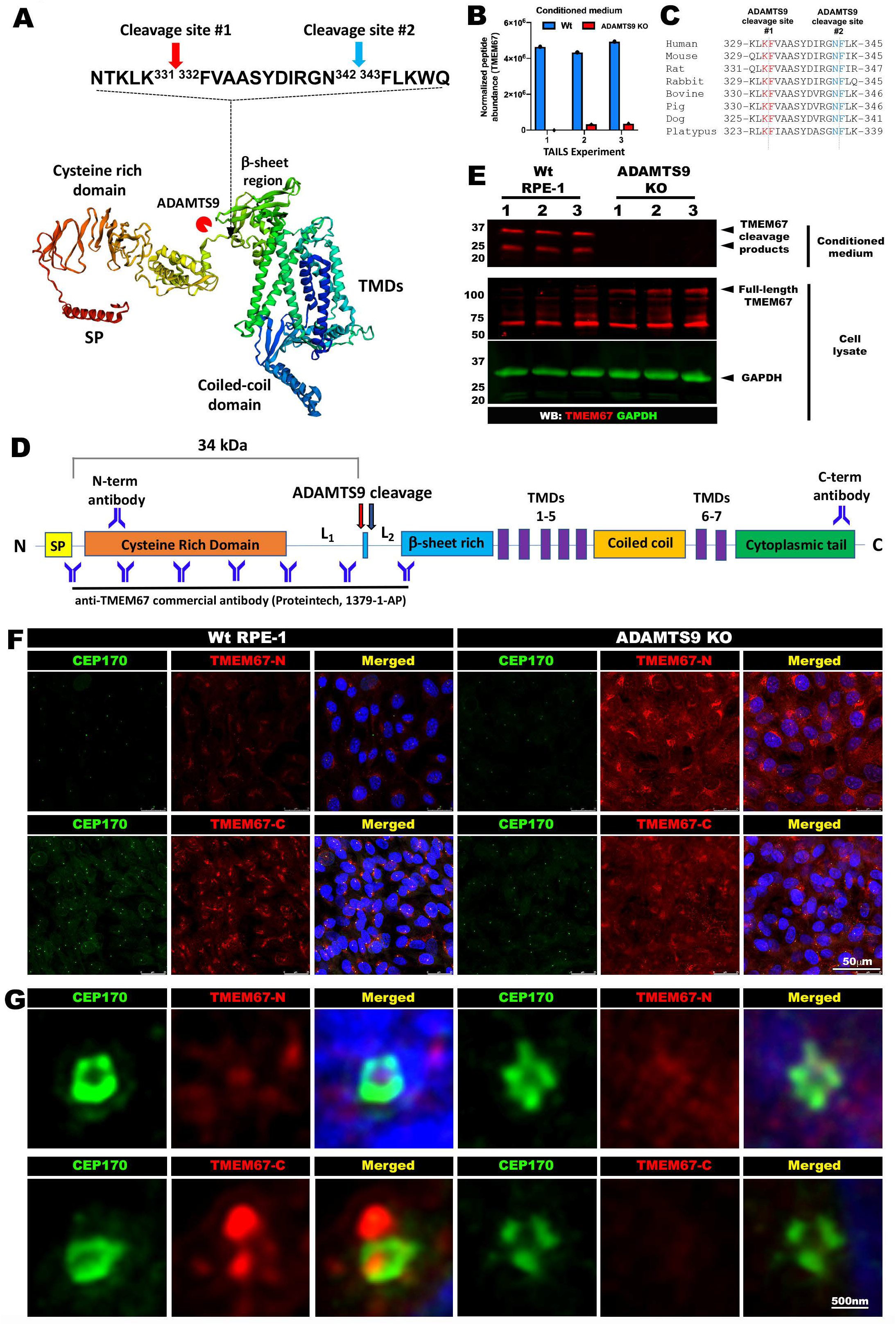
Identification and validation of TMEM67 as a novel substrate of ADAMTS9. (**A**) AlfaFold model of TMEM67 indicating the two novel ADAMTS9 cleavage sites identified utilizing N-terminomics, which are in a linker region in between the Cysteine rich domain (CRD) and the β-sheet rich domain (BRD) of TMEM67. (**B**) TMEM67 ^332^FVAASYDRGN^342^ peptide abundance from triplicate TAILS experiments comparing the conditioned medium of wild type (blue) and *ADAMTS9* KO (red) RPE-1 cells. (**C**) TMEM67 amino acids sequence alignment showing conservation of the identified cleavage residues and the surrounding amino acids throughout mammalians. (**D**) Updated diagram of TMEM67 domain structure, indicating the two ADAMTS9 cleavage sites, the newly identified linkers 1 and 2, the binding sites of N-terminal or C-terminal specific TMEM67 antibodies or the TMEM67 commercial antibody which spans the cleavage site. (**E**) Western blot showing two closely migrating TMEM67 bands matching the predicted molecular weight of 34 kDa by ADAMTS9-mediated N-terminal cleavage, present in the wild type conditioned medium but absent in the *ADAMTS9* KO RPE-1 conditioned medium. The corresponding cell layers show more full-length TMEM67 abundant in the *ADAMTS9* KO RPE-1 cells compared to wild type. (**F-G**) Conventional high resolution confocal microscopy (**F**), or super resolution confocal microscopy (**G**), showing co-immunostaining of TMEM67 N-terminal or C-terminal specific antibodies (red) with the mature basal body marker CEP170 (green) in serum starved RPE-1 cells. Only the TMEM67 C-terminus is colocalized with CEP170 in wild type RPE-1 cells, which is lost in *ADAMTS9* KO RPE-1 cells which shows increased N-terminal and C-terminal staining throughout the cell. Arrowhead in **G** indicates the presumed transition zone localization of the TMEM67 C-terminal fragment. Scale bar in **F** is 50μm and 500nm in **G**.

### Removal of the TMEM67 N-terminus is required for ciliogenesis

To investigate the requirement for TMEM67 cleavage on ciliogenesis, we generated mammalian expression constructs for full-length TMEM67 (TMEM67-FL), the C-terminal half resulting from ADAMTS9 cleavage (TMEM67 Δ342), the N-terminal CRD fragment shed by ADAMTS9 cleavage (TMEM67 N-331), and non-cleavable TMEM67, in which both (S1+S2) or individual cleavage sites were mutated into alanine residues (**Fig. 2A**). Importantly, TMEM67 Δ342 was cloned into pSecTag2c, adding a signal peptide N-terminal to F^343^ to retain the correct topology of the cleaved TMEM67 C-terminal fragment. Loss of ciliogenesis in *TMEM67* KO RPE-1 cells was rescued by the transfection of full length TMEM67 or TMEM67 Δ342 but not by TMEM67-N-331 or by the cleavage site mutants (S1, S2, S1+S2) (**Fig. 2B-C**). Loss of ciliogenesis in *ADAMTS9*-null RPE- 1 cells could also be partially restored by introducing TMEM67 Δ342 (**Fig. 2D-E**). These data together indicate that TMEM67 cleavage is required for ciliogenesis and that the TMEM67 C-terminal half (TMEM67 Δ342) generated by ADAMTS9-mediated cleavage is sufficient and necessary to restore ciliogenesis. We next asked whether TMEM67 cleavage is required for Its TZ localization. Full-length TMEM67 and TMEM67 Δ342 exhibited TZ localization comparable to that of Wt cells, while TMEM67 N-331, or the cleavage site mutants (S1, S2 and S1+S2) showed no TZ localization (**Fig. 2F**). These data show that TMEM67 cleavage regulates its C-terminal localization to the TZ and ciliogenesis.

**Figure 2:**
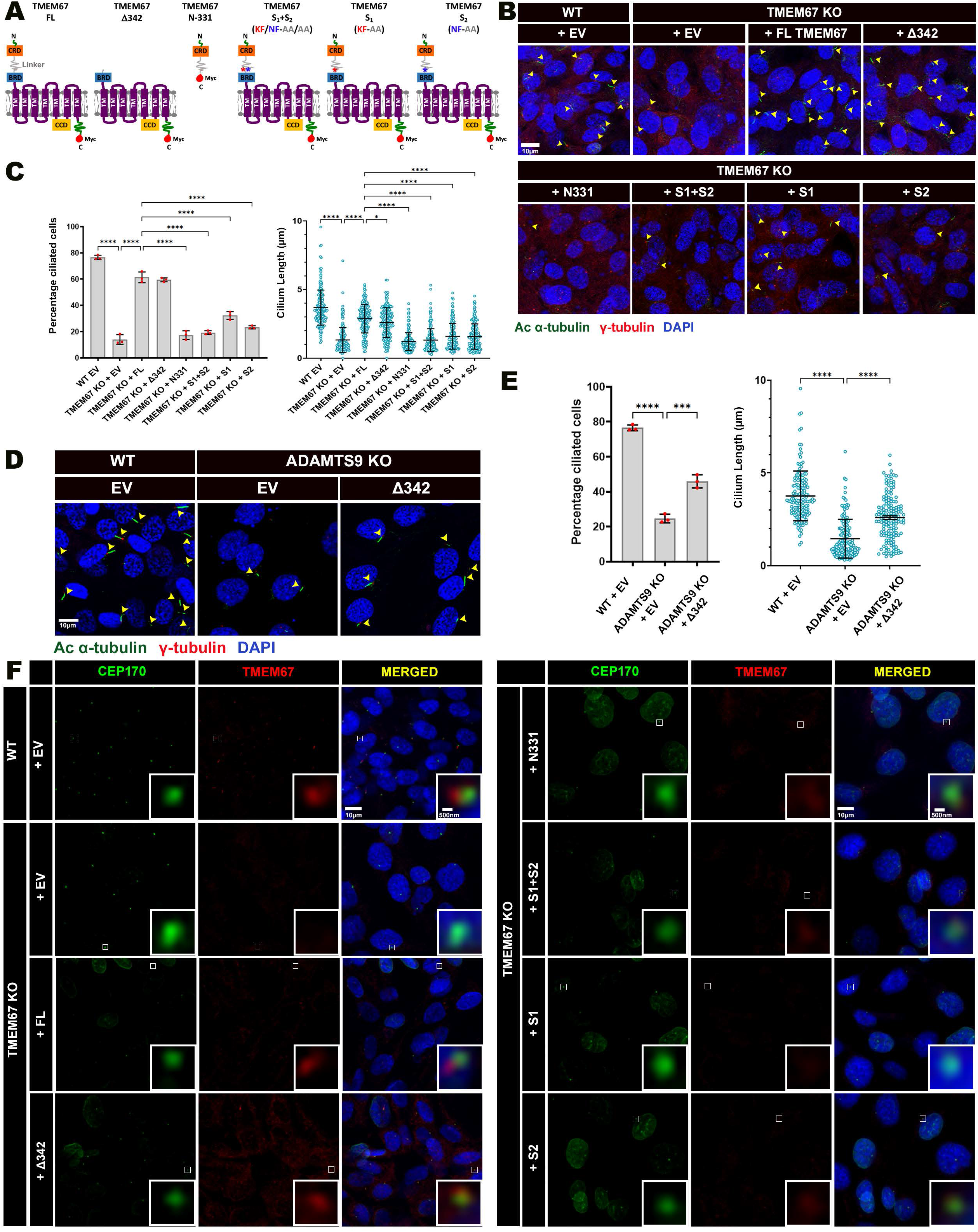
ADAMTS9-mediated TMEM67 cleavage is required for ciliogenesis and TMEM67 TZ localization. (**A**) TMEM67 constructs generated encoding full length TMEM67 (FL), the C-terminal cleavage product (Δ342), N-terminal cleavage product (N-331), dual cleavage mutant (S1+S2), individual mutation of cleavage site 1 (S1) and cleavage site 2 (S2) alone. A signal peptide sequence was added at the N-terminus of the TMEM67 Δ342 construct for the correct synthesis of the molecule with an N-terminal extracellular domain, and is not indicated in the illustration. (**B-C**) TMEM67 KO RPE-1 cells transfected with full-length TMEM67 or the TMEM67 Δ342 can rescue ciliogenesis but TMEM67 N-331 or cleavage site mutants do not. Yellow arrowheads indicate primary cilia marked by acetylated α-tubulin (green) and the basal bodies marked by γ- tubulin (red). Percentage of ciliated cells and cilium length quantification from 3 independent experiments are shown in **C**. (**D-E**) TMEM67 Δ342 can partially rescue ciliogenesis and cilium length of *ADAMTS9* KO RPE-1 cells. Yellow arrowheads indicate primary cilia marked by acetylated α-tubulin (green) and the basal bodies marked by γ-tubulin (red). Percentage of ciliated cells and cilium length quantification from 3 independent experiments are shown in **E.** **** indicates a p-value < 0.0001, *** < 0.001, ** < 0.01, * <0.05 in Mann-Whitney U test for statistical significance. Error bars indicate Mean ± S.D. (F) TMEM67 (red) and CEP170 (green) immunostaining in Wt and *TMEM67* KO cells transfected with TMEM67 constructs show transition zone localization of TMEM67 FL and TMEM67 Δ342 constructs only. Scale bars in **B**, **D** and **F** are 10μm and 500nm in the **F** inserts. **** indicates a p- value < 0.0001, *** < 0.001, ** < 0.01, * <0.05 in One-way ANOVA test for statistical significance. Error bars indicate Mean ± S.D.

### TMEM67 cleavage is required for the assembly of the MKS/B9 module of the ciliary transition zone

To gain further insight into the molecular mechanism of how loss of TMEM67 cleavage affects ciliogenesis, we investigated how TMEM67 and ADAMTS9 loss affected TZ assembly. We performed a comprehensive immunostaining analysis of 14 TZ proteins in Wt, *TMEM67* KO and *ADAMTS9* KO RPE-1 cells, utilizing high-resolution and super-resolution confocal microscopy to investigate their localization to the matured basal body upon ciliogenesis induction. Of the 14 TZ proteins examined, we found 6 proteins (TCTN1, TCTN2, TCTN3, TMEM237, CC2D2A and B9D2) were significantly reduced in both *TMEM67* and *ADAMTS9* KO cells (**Fig. 3A-B**, **Sup.** Figures 3-4). In addition to these core changes, loss of ADMTS9 but not TMEM67 significantly reduced B9D1, NPHP1, and NPHP5 and significantly increased CEP290 staining, while loss of TMEM67 but not ADAMTS9 significantly reduced RPGRIP1L and INPP5E localization to the mature ciliary base (**Fig. 3B**). Intensity of the sub-distal appendage protein CEP170 was measured as a control and showed no significant change between the cell lines (**Fig. 3B**). We carried out western blotting and qRT-PCR analysis of the 6 MKS/B9 module proteins that were reduced in both cell lines, which showed total protein levels were also reduced (**Fig. 3C**), but their gene expression was not significantly decreased in TMEM67 and ADAMTS9 KO cells compared to Wt cells, except *TCTN3* and *CC2D2A,* which showed reduced transcription in both cell lines (**Fig. 3D**). These results, summarized in the known TMEM67 interactome (**Fig. 3E**), shows loss of TMEM67 and ADAMTS9 proteolytic activity significantly affects the MKS/B9 module during TZ assembly.

**Figure 3:**
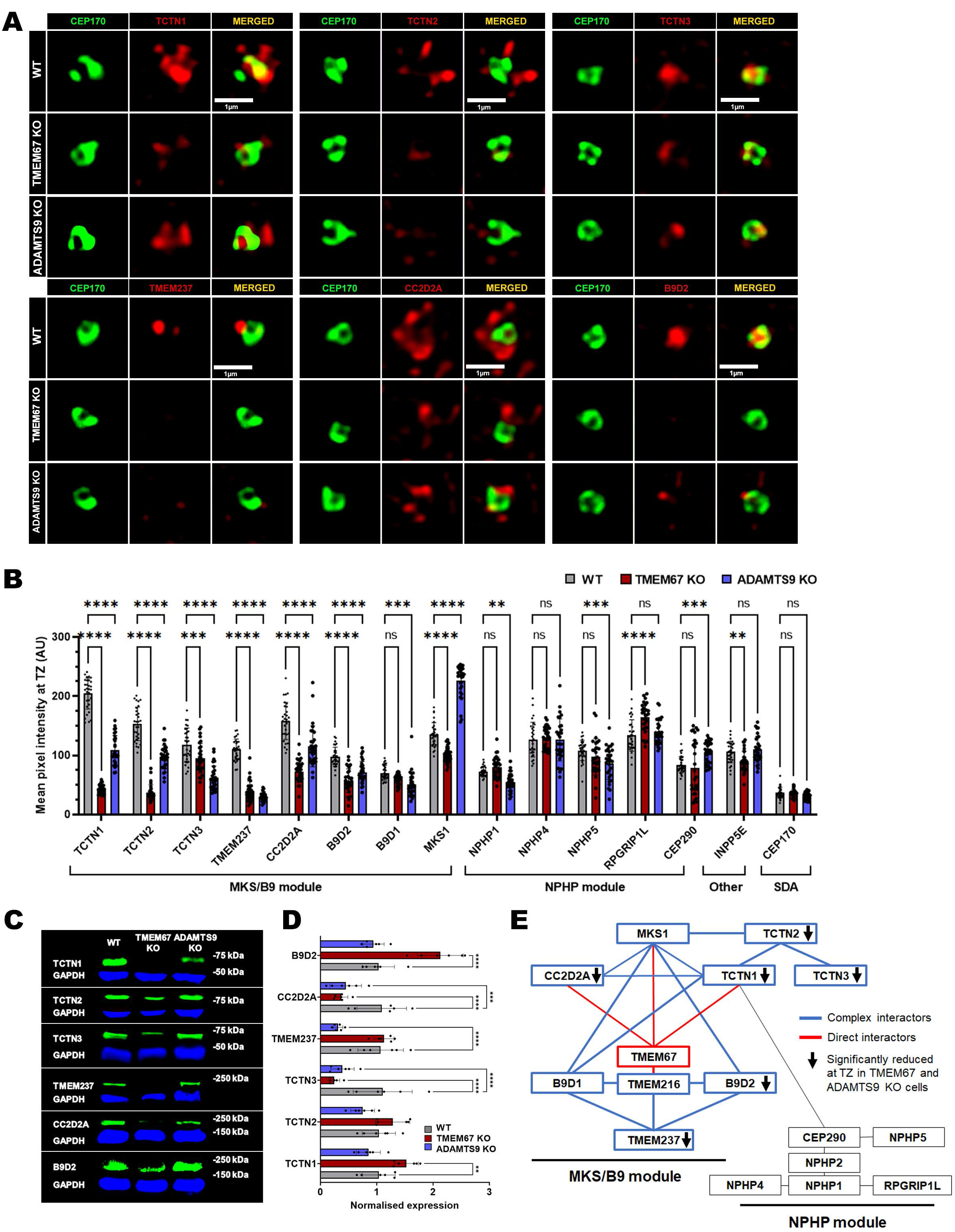
Loss of TMEM67 or ADAMTS9 severely impairs transition zone assembly in RPE- 1 cells. (**A**) Super-resolution confocal microscopy images of the transition zone MKS/B9 module proteins TCTNT1, TCTN2, TCTN3, TMEM237, CC2D2A and B9D2 (red), co-stained with the sub-distal appendage protein CEP170 (green), show significantly decreased or loss of localization at the mature basal bodies of *ADAMTS9* KO and *TMEM67* KO RPE-1 cells in comparison to wild type. (**B**) Mean pixel intensity quantification at the ciliary base of all 14 transition zone proteins analyzed in this study comparing serum starved wild type, *TMEM67* KO and *ADAMTS9* KO RPE-1cells (n= 50, each group). **** indicates a p-value < 0.0001, *** < 0.001, ** < 0.01, * <0.05 in one-way ANOVA test for statistical significance. Error bars indicate Mean ± S.D. (**C**) Western blot analysis of the 6 transition zone proteins TCTN1, TCTN2, TCTN3, TMEM237, CC2D2A and B9D2 (green) show reduced total protein levels in *TMEM67* KO cells and unaltered levels in *ADAMTS9* KO cells. (**D**) qRT-PCR analysis for *TCTN1*, *TCTN2*, *TCTN3*, *TMEM23*7, *CC2D2A* and *B9D2* transcript levels show similar or significantly increased transcription for *TCTN1*, *TCTN2* and *B9D2* while *TCTN3* and *CC2DA* show decreased expression levels in TMEM67 and ADAMTS9 KO cells in comparison to wild type cells while *TMEM237* expression is decreased in ADAMTS9 KO cells and unchanged in *TMEM67* KO cells. **** indicates a p-value < 0.0001, *** < 0.001, ** < 0.01, * <0.05 in one-way ANOVA test for statistical significance. Error bars indicate Mean ± S.D. (**E**) Graphical representation of the transition zone protein interaction network in relation to TMEM67 (red box). The TMEM67 direct interactions are shown in red lines while interactions amongst the four direct interactors are shown in blue lines. The MKS/B9 module proteins are shown in blue boxes while the NPHP module proteins are shown in black boxes. Black arrows indicate significantly reduced TZ proteins in both *TMEM67* KO and *ADAMTS9* KO cells. Scale bar in **A** is 500nm.

### The TMEM67 C-terminal half produced by ADAMTS9-mediated cleavage (TMEM67 Δ342) is sufficient for restoring ciliary transition zone assembly

Since loss of ciliogenesis in *TMEM67* KO and *ADAMTS9* KO cells can be rescued by TMEM67 Δ342, we tested whether it was sufficient to rescue the loss of the TZ MKS/B9 module. *TMEM67* KO cells transfected with TMEM67 Δ342 showed significantly increased levels of the 6 MKS/B9 proteins reduced from the TZs lacking TMEM67 (**Fig. 4A-B**). *ADAMTS9* KO cells transfected with TMEM67 Δ342 showed a similar result with significantly increased TZ pixel intensity for TCTN1, TCTN2, TCTN3, TMEM237, and CC2D2A with the exception of B9D2 (**Fig. 4C**). These results show that TMEM67 Δ342 is sufficient to rescue TZ assembly defects caused by loss of TMEM67 and in part in *ADAMTS9* deficient cells. This suggests that release of the TMEM67 Δ342 fragment following ADAMTS9-mediated cleavage is an important regulatory mechanism governing the recruitment or stabilization of membrane associated MKS/B9 components to the TZ. This function is most likely regulated by the intact cytoplasmic coiled-coil domain (CCD), present in the TMEM67 C-terminus, and many other ciliary proteins, which utilize them for dimerization and protein interactions [40–45].

**Figure 4:**
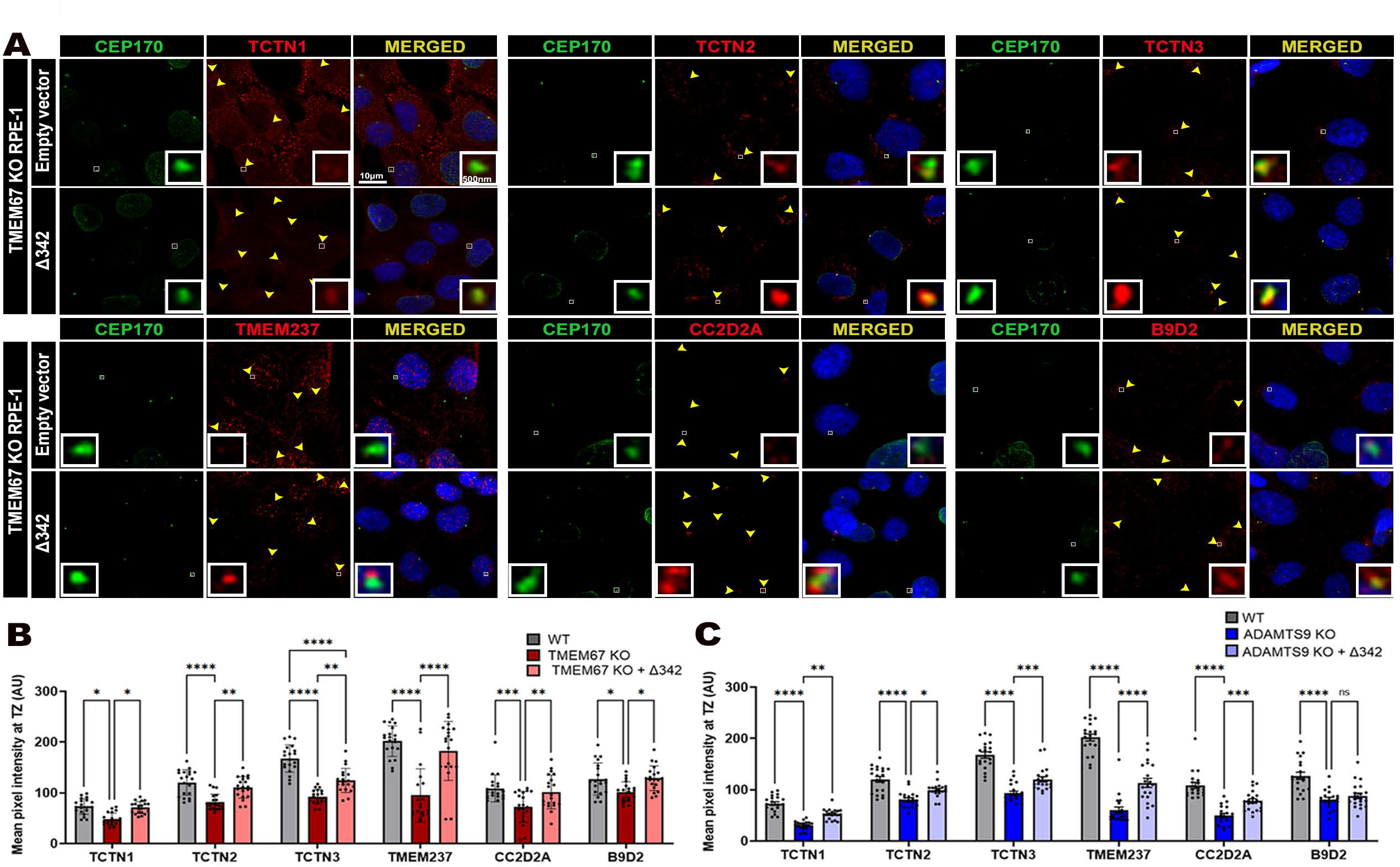
The TMEM67 C-terminal cleavage fragment can restore TZ assembly of *TMEM67* KO and *ADAMTS9* KO cells. (**A**) Immunostaining of the 6 ciliary transition zone proteins affected by TMEM67 loss (red) and the sub-distal appendage protein CEP170 (green) in *TMEM67* KO RPE-1 cells transfected with empty vector (EV) or the TMEM67 C-terminal cleavage product Δ342 show restoration of TZ assembly. Transition zone/ CEP170 foci are indicated by yellow arrowheads and are shown in high magnification in the inserts. (**B-C**) Transition zone mean pixel intensity quantification of the six transition zone proteins analyzed shows statistically significant rescue of nearly all six TZ proteins in *TMEM67* KO and *ADAMTS9* KO cells upon TMEM67 Δ342 transfection. Scale bar in **A** is 10µm and 500nm in the inset. **** indicates a p-value < 0.0001, *** p-value < 0.001, ** p-value < 0.01, * p-value < 0.05 in two-way ANOVA for statistical analysis. Error bars indicate Mean ± S.D.

### *TMEM67* variants in the cleavage site or adjacent residues result in ciliopathies

Three *TMEM67* patient mutations within or adjacent to the TMEM67 cleavage motif have been identified in ciliopathy patients (**Fig.5A**). The c.1027T>G mutation corresponding to p.F343V, occurs at the second cleavage site and resulted in bilateral enlarged cystic kidneys, ductal plate malformation (DPM) of the liver and abnormal lung lobulation [46]. A c.986A>C mutation leading to p.K329T, two residues upstream of the first cleavage site, causes nephronophthisis (NPHP), ataxia, cerebellar vermis hypoplasia, mental retardation, and hepatic fibrosis [19]. The c.1046T>C mutation, corresponding to p.L349S, 6 residues downstream of the second cleavage site was identified in individuals across two families with biallelic *TMEM67* mutations, resulted in MKS, COACH syndrome, cleft palate and intra-uterine growth retardation [17]. We tested whether these *TMEM67* patient variants affect ciliogenesis by conducting rescue experiments in *TMEM67* KO RPE-1 cells. RPE-1 cells transfected withTMEM67 F343V, K329T and L349S variants showed reduced ciliogenesis compared to *TMEM67* KO cells transfected with full-length TMEM67 (**Fig. 5B-C**). All three patient variants also showed reduced TZ localization comparable to that of non-cleavable TMEM67 constructs S1 + S2, S1 and S2 (**Fig. 5D**). Interestingly, all three variants are in the two novel linker regions identified here in between the CRD and BRD domains of TMEM67. These results show that the cleavage residues and the correct formation of the two linker regions are critical to TMEM67 functionality during human development.

**Figure 5:**
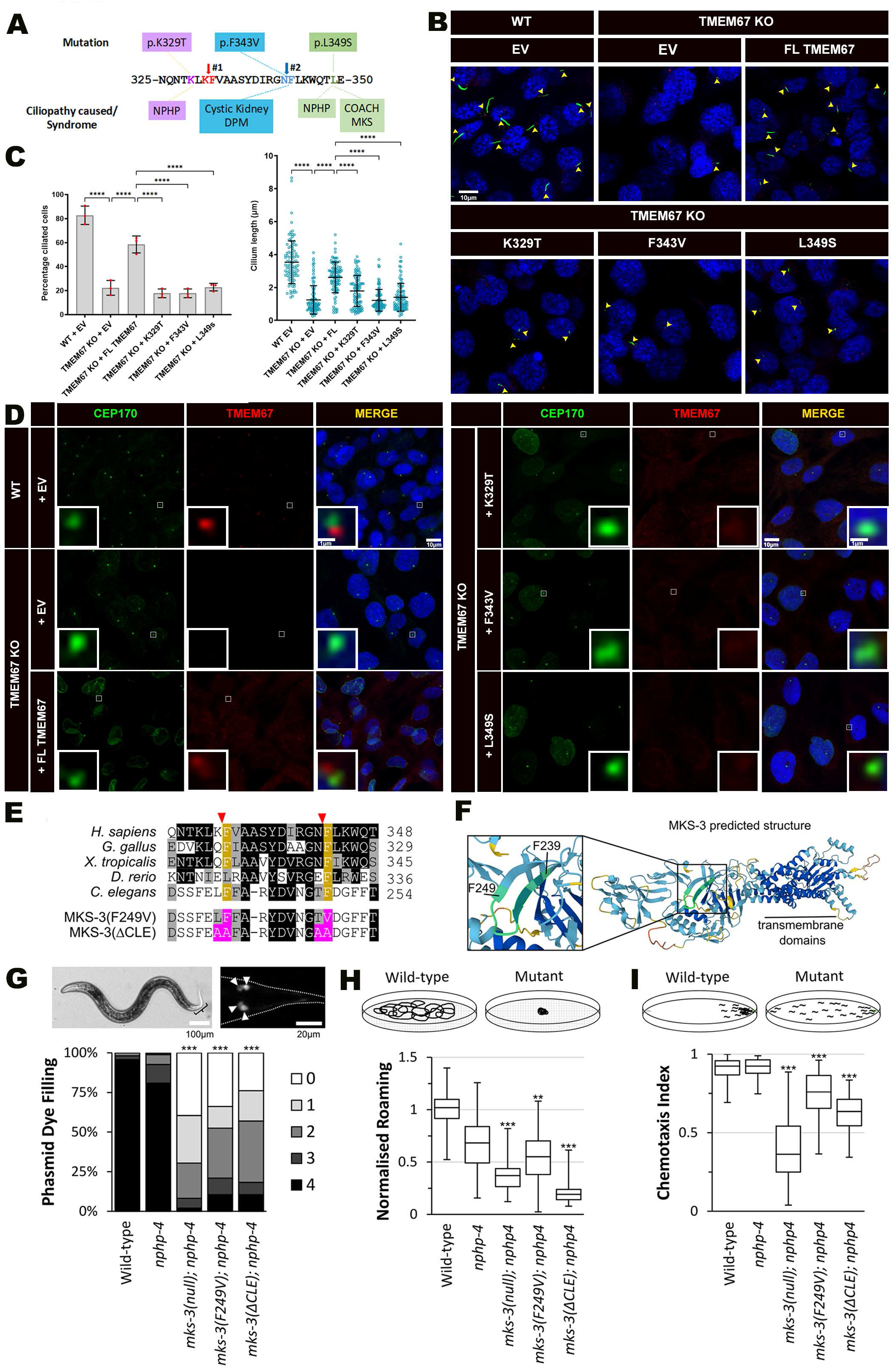
Ciliopathy patient variants within or surrounding the TMEM67 cleavage residues affect normal ciliogenesis and essential for mks-3 function in *C elegans*. (**A**) *TMEM67* patient mutations K329T, F342V and L349S in relation to the cleavage residues and the underlying ciliopathies identified in each variant. (**B-C**) *TMEM67* KO cells transfected with the patient variants K329T, F342V and L349S fail to rescue ciliogenesis and cilium length of serum starved RPE-1 cells. Yellow arrowheads indicate primary cilia immunolabelled with acetylated α-tubulin (green) and γ-tubulin (red). Percentage of ciliated cells and cilium length from three independent experiments are shown in **C**. **** indicates a p-value < 0.0001, *** < 0.001, ** < 0.01, * <0.05 in one-way ANOVA test for statistical significance. Error bars indicate Mean ± S.D. (**D**) Immunolabelling of wild type and *TMEM67* KO RPE-1 cells with TMEM67 (red) and CEP170 (green) after transfection with the indicated constructs showing the patient variant forms K329T, F342V and L349S fail to localize to the mature basal body. (**E**) Amino acid alignment of the two TMEM67 cleavage sites (red triangles) and the surrounding residues in human, chicken, frog, zebrafish and worm show high degree of conservation during evolution. Identical amino acids are highlighted in black and similar amino acids are highlighted in grey. Conservation of the phenylalanine residues common to both cleavage sites are shown by the yellow highlights. Amino acid changes made in the MKS-3 F249V form (modeling the ciliopathy patient variant F343V of the 2^nd^ cleavage site) and the MKS-3 ΔCLE form where both cleavage site residues are changed to alanine are shown in the bottom. (**F**) AlphaFold modeled structure of *C. elegans* MKS-3 (AF-Q20046-F1) showing the cleavage sites. Color of the backbone indicates AlphaFold confidence score (dark blue = very high, blue = high, yellow = low, orange = very low). The inset shows residues F239-F249 (highlighted in green) with the boxed area shown in higher magnification. (**G**) Assessment of cilium integrity using a lipophilic dye (DiI/DiO) uptake assay shows significantly reduced phasmid dye filling in the two cleavage site mutant worm lines are comparable to the mks-3 null worms. Fluorescence image shows the four ciliated neurons (phasmids) in the worm’s tail that incorporate DiI/DiO. Arrowheads indicate the phasmid cell bodies. Histogram shows the percentages of worms that incorporate dye in 0-4 phasmid neurons. (**H**) Assessment of sensory cilium function using a single worm assay of cilia-dependent foraging behavior show significantly lower roaming in the MKS-3 cleavage site mutant worms normalized to wild type level. (**I**) Assessment of sensory cilium function using a population-based assay of chemotaxis towards the chemical attractant benzaldehyde show significantly lower sensory cilium functionality in MKS- 3 cleavage site mutant worms. Box plots indicate the maximum and minimum values (bars), median, lower quartile, and upper quartile. *** indicates a p-value < 0.001 and ** a p-value < 0.01. Scale bars in **B** and **D** are 10μm and 1μm in the **D** inserts, 100µm and 20µm in **G**.

### The *TMEM67* cleavage site is essential for the functionality of the *C. elegans* ortholog (MKS-3)

Using the ConVarT sequence alignment tool [47], we found that the predicted cleavage sites occur within a highly conserved region of TMEM67 orthologs across diverse animal species, including the small roundworm, *Caenorhabditis elegans* (**Fig. 5E**). AlphaFold [48] analysis revealed that this region of the *C. elegans* ortholog (MKS-3) is on an outer face of the protein, consistent with its accessibility for extracellular protease cleavage (**Fig. 5F**). To determine if the predicted cleavage site is essential for MKS-3 function, CRISPR-Cas9 methodology was used to engineer two variants: F249V that corresponds to the F343V patient variant allele of the second cleavage site, and mks-3(ΔCLE), which possesses four amino acid substitutions (L238A, F239A, T248A, F249A) predicted to abolish both cleavage sites similar to the human TMEM67 S1+S2 construct. To assess MKS-3 function, quantitative assays of sensory cilia integrity (dye filling) and function (roaming, chemotaxis) were employed [28]. As *mks-3* functions redundantly with NPHP genes to control cilium formation and function in worms, the *mks-3* alleles were assessed in a sensitized *nphp-4* mutant background [49, 50]. In all three assays, the F249V and ΔCLE mutations significantly disrupted MKS-3 function (**Fig. 5G-I**). Indeed, the cilium integrity defect (dye filling) approached that of the *mks-3* null allele, the cilium function abnormalities were somewhat less severe. Taken together, these data show that human TMEM67 cleavage sites are essential for *C. elegans* MKS-3 ciliary functions. Whether the ancestral nematode ortholog of the mammalian ADAMTS9, GON-1 [51], is also involved in MKS-3 cleavage remains to be determined.

### *Tmem67^ΔCLE/ΔCLE^* mice develop severe cystic kidneys and hydrocephalus and phenocopy *Tmem67*-null mice

To investigate the role of TMEM67 cleavage *in vivo*, we developed a mouse model in which both the TMEM67 cleavage sites K^331^/F^332^ (cleavage site 1) and N^342^/F^343^ (cleavage site 2) were mutated to alanine residues (*Tmem67^ΔCLE/ΔCLE^*, **Sup.** Fig. 5A). Two independent founder lines, *Tmem67^ΔCLE1^* and *Tmem67^ΔCLE2^*, were generated from two independent mouse ES cell clones in the C57BL6/j background (**Sup.** Fig. 5A). Homozygous mice from both lines were phenotypically identical and here forth will be commonly referred to as *Tmem67^ΔCLE/ΔCLE^* mice. *Tmem67^ΔCLE/ΔCLE^* mice only survived till post-natal day 14 (p14) and phenocopied *Tmem67-null* mice, developing large polycystic kidneys and hydrocephaly with significantly impaired postnatal growth (**Fig. 6A-C**, **Sup. Fig. 6**, **Sup.** Fig. 8A-C). The *Tmem67^ΔCLE/+^* (heterozygous) mice were phenotypically normal and showed normal kidney histology at 10 weeks of age (p70, **Sup. Fig-7A**). The vast majority of *Tmem67^ΔCLE/ΔCLE^* mice exhibited embryonic lethality similar to *Tmem67* KO mice [27], and showed defective cardiac, hepatic and vascular phenotypes (**Sup.** Fig. 7B-C). *Tmem67^ΔCLE/ΔCLE^*hearts from E15.5 and E18.5 embryos revealed severely impaired cardiac development, resulting in an overriding-aorta, ventricular septal defect (VSD), and a loss of myocardial compaction (**Sup.** Fig. 7C). Livers from postnatal *Tmem67^ΔCLE/ΔCLE^* mice showed defective hepatic portal vein (HPV) branching morphogenesis, loss of hepatocyte differentiation, and increased fibrosis (**Sup.** Fig. 8E-G). Scanning electron microscopy (SEM) revealed *Tmem67^ΔCLE/ΔCLE^* cystic renal tubular epithelium had severely shortened, malformed primary cilia (turquoise arrows), as well as extended primary cilia which were morphologically abnormal (orange arrows), similar to that seen in *Tmem67-*null kidneys (**Fig. 6D**). Transmission electron microscopy (TEM) of both short and long cilia types showed a complete loss of the TZ necklace formation, seen in the Wt littermate kidney primary cilia (red arrow heads, **Fig. 6E**). Loss of TMEM67 results in hydrocephaly in humans, mice, rat, and zebrafish [26, 52–54]. Analysis of the multiciliated ependymal epithelium lining the lateral brain ventricles showed highly abnormal motile cilia formation in the *Tmem67^ΔCLE/ΔCLE^* mice (**Fig. 6F**). Nearly all the motile cilia observed contained bulbous tips and large ciliary membrane bulges either at the distal ends or on their sides (yellow arrowheads, **Fig. 6F**). Compared to the well-organized motile cilia observed in Wt brain ventricles, the ependymal cilia of the *Tmem67^ΔCLE/ΔCLE^* brain ventricles were tangled and laid flat on the ependymal cell surface (**Fig. 6F**). TEM analysis of the abnormal motile cilia of the *Tmem67^ΔCLE/ΔCLE^* brain ventricles revealed a phenotype similar to the renal primary cilia and showed a complete loss of the TZ necklace formation (red arrowheads, **Fig. 6G**). These results validated our *in vitro* findings analyzing *TMEM67* and *ADAMTS9*-null RPE-1 cells. These data also show that TMEM67 cleavage is essential for normal mammalian development and the normal morphogenesis of the TZ of both primary and motile cilia, and its loss leads to multi-organ failure in mice. Crucially, since *Tmem67^ΔCLE/ΔCLE^* mice are phenotypically identical to *Tmem67*-null mice, it also suggests that TMEM67 loss-of-cleavage results in a nonfunctional TMEM67 in ciliogenesis.

**Figure 6:**
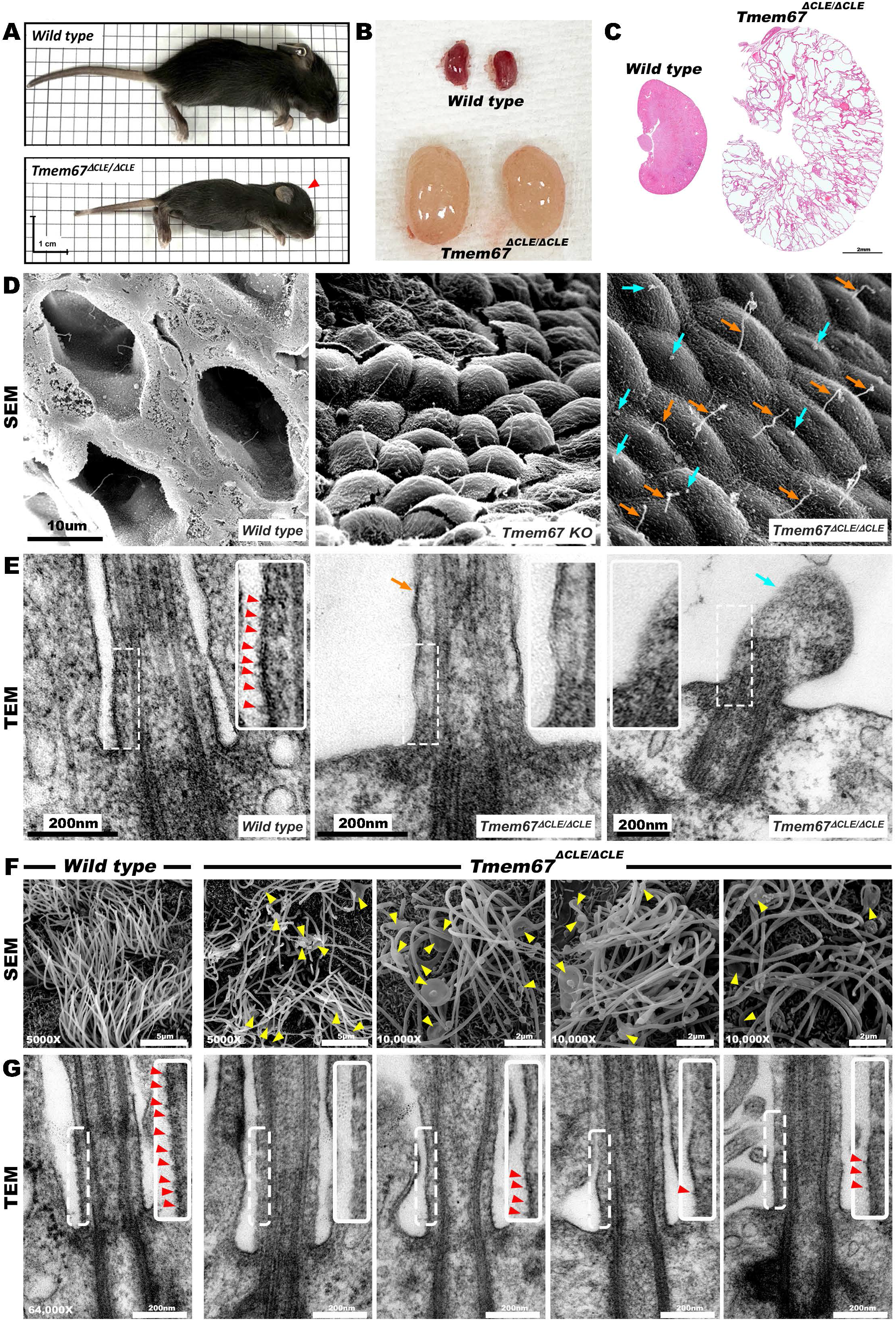
Characterization of a novel *Tmem67^ΔCLE/ΔCLE^* mouse model reveal loss of TMEM67 cleavage leads to ciliopathy formation in mammals. (**A**) *Wild type* and *Tmem67^ΔCLE/ΔCLE^* littermates at postnatal day 14 (p14) showing impaired growth and a dome shaped head (red arrowhead) indicative of hydrocephalus formation in *Tmem67^ΔCLE/ΔCLE^* mice. (**B**) Kidneys dissected from *wild type* and *Tmem67^ΔCLE/ΔCLE^* littermates at p14 showing highly enlarged polycystic kidneys in the mutant mice. (**C**) Hematoxylin and Eosin stained kidney sections from p14 mice show highly cystic renal histopathology in the *Tmem67^ΔCLE/ΔCLE^* mutant kidneys compared to that of a *wild type* littermate. (**D**) Freeze-fracture scanning electron microscopy (SEM) images show defective primary cilia formation in *Tmem67^ΔCLE/ΔCLE^* kidneys similar to *Tmem67* KO kidneys. The turquoise arrows indicate extremely short cilia and orange arrows indicate extended primary cilia with highly abnormal morphologies present on the *Tmem67^ΔCLE/ΔCLE^* mutant cystic renal tubular epithelium. (**E**) Transmission electron microscopy (TEM) images of primary cilia showing the transition zone “necklace” (red arrowheads) in a *wild type* cilium is entirely missing in both types of *Tmem67^ΔCLE/ΔCLE^*primary cilia. The turquoise and orange arrows indicate extremely short or an extended primary cilium section visualized by TEM. (**F**) SEM images showing highly abnormal, tangled motile cilia with large membrane bulges (yellow arrowheads) and bulbus tips present in the multiciliated ependymal cells lining the lateral brain ventricles in *Tmem67^ΔCLE/ΔCLE^* mice compared to the uniformly organized motile cilia seen in the *wild type* brain ventricles. (**G**) TEM images show a similar loss of the transition zone necklace formation (red arrowheads) in the *Tmem67^ΔCLE/ΔCLE^*motile ependymal cilia in comparison to *wild type* ependymal cilia. Scale bar in **C** is 2mm, 10µm in **D,** 5µm or 2 µm in **F** and 200nm in **E** and **G**.

### Non-cleavable TMEM67 maintains the ability to transduce both canonical and non-canonical Wnt signaling but is deficient for ciliogenesis and TZ localization

To investigate the functionality of the TMEM67- ΔCLE protein in more detail, we harvested and immortalized mouse embryonic fibroblasts (MEFs) from E13.5 *Tmem67^ΔCLE/ΔCLE^*embryos. Following serum starvation, *Tmem67^ΔCLE/ΔCLE^* MEFs had significantly reduced ciliogenesis and cilia lengths compared to *Tmem67^ΔCLE/+^* MEFs (**Fig. 7A-B**). TZ localization of TMEM67 was also lost in *Tmem67^ΔCLE/ΔCLE^* MEFs (**Fig. 7C**), supporting the observation that the human TMEM67- S1+S2 non-cleavable protein did not rescue ciliogenesis, and showed lack of TZ localization in RPE-1 cells (**Fig. 2F**). Transfection of the TMEM67 C-terminal half (TMEM67 Δ342), significantly improved the loss of ciliogenesis and cilia lengths of *Tmem67^ΔCLE/ΔCLE^* MEFs (**Fig. 7D-E**). Western blot analysis of the serum free conditioned media revealed a ∼34kDa TMEM67cleavage product produced by the Wt MEFs but not by the *Tmem67^ΔCLE/ΔCLE^* MEFs, with more full-length TMEM67 in the *Tmem67^ΔCLE/ΔCLE^* cell lysates (**Fig. 7F**). qRT-PCR analysis for *Tmem67* transcript levels showed similar abundance in MEFs (**Fig. 7G**) and whole kidneys (**Sup.** Fig. 5C), which show *Tmem67* transcription is unaffected in *Tmem67^ΔCLE/ΔCLE^* mice. Combined, these data validated the novel *Tmem67^ΔCLE/ΔCLE^* mouse model’s ability to produce a non-cleavable form of TMEM67, that lacks ciliary TZ localization and TZ assembly.

**Figure 7:**
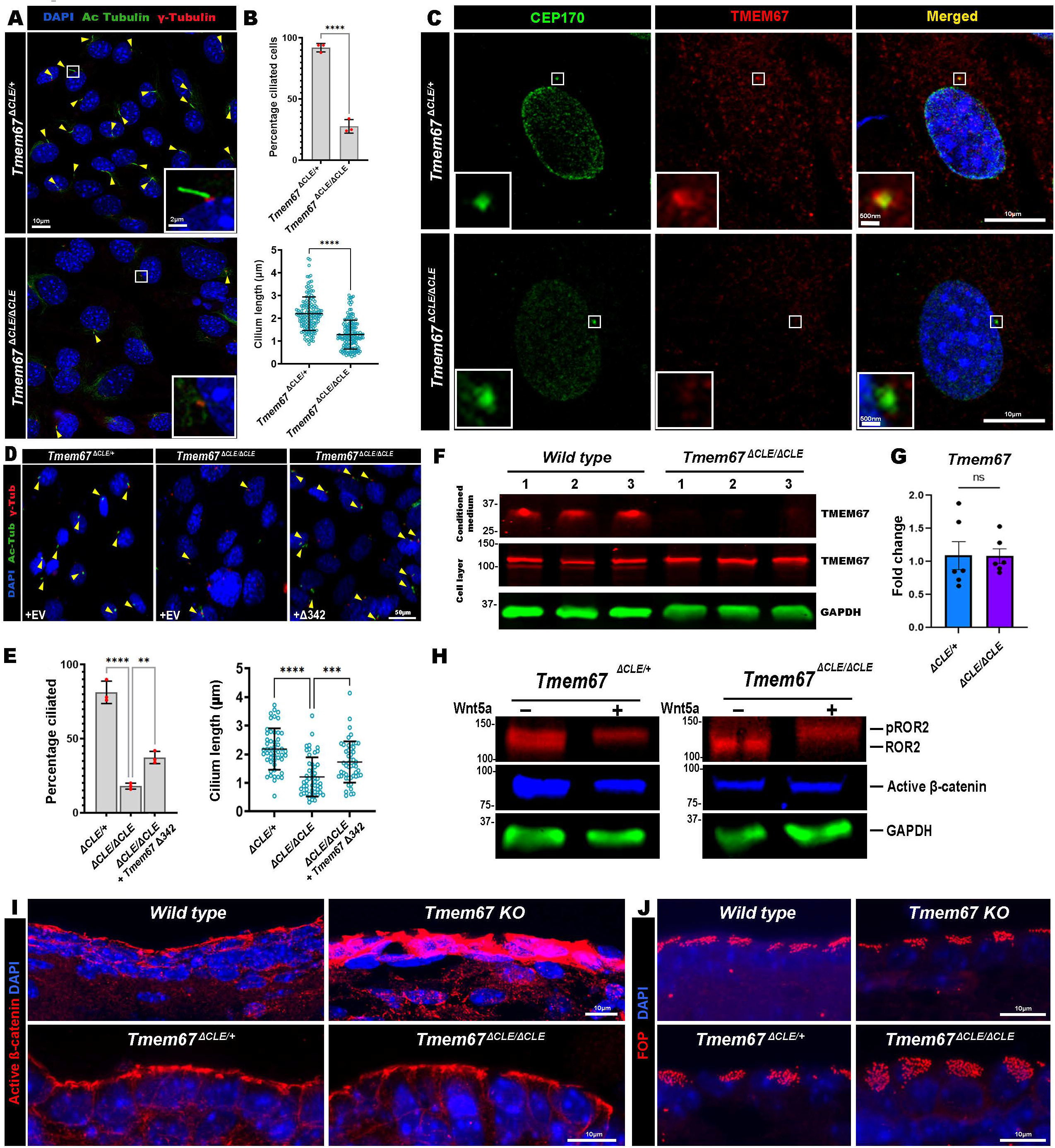
Loss of ciliogenesis but normal Wnt Signaling in *Tmem67^ΔCLE/ΔCLE^* embryonic fibroblasts and brains. (**A-B**) Serum starved *Tmem67^ΔCLE/ΔCLE^* mouse embryonic fibroblasts (MEFs) show significantly shorter and fewer primary cilia (yellow arrowheads) marked by acetylated α-tubulin (green) and γ-tubulin (red) staining. (**C**) CEP170 (green) and TMEM67 (red) immunostaining of serum starved *Tmem67^ΔCLE/ΔCLE^* MEFs show loss of TZ localization of non-cleavable TMEM67. (**D-E**) Transfection of *Tmem67^ΔCLE/ΔCLE^* MEFs with TMEM67 1342 significantly improves ciliogenesis and cilium length. (**F**) Western blotting for TMEM67 (red bands) and GAPDH (green bands) show presence of the 34kDa TMEM67 N-terminal fragment in the *wild type* MEF conditioned medium and its absence in the *Tmem67^ΔCLE/ΔCLE^*MEF conditioned medium from triplicate cultures from each group. The corresponding cell layers show increased full-length TMEM67 levels in the *Tmem67^ΔCLE/ΔCLE^* MEF cell lysates. (**G**) qRT-PCR analysis show similar levels of *Tmem67* transcript in the *Tmem67^ΔCLE/+^* and *Tmem67^ΔCLE/ΔCLE^* MEFs. (**H**) Western blot showing ROR2 (red) phosphorylation is unaffected and *Tmem67^ΔCLE/ΔCLE^* MEFs and undergo normal non-canonical Wnt signaling in response to Wnt5a treatment. Blue, active Δ- catenin, Green, GAPDH. (**I**) Active Δ-catenin immunostaining (red) of *Wt*, *Tmem67 KO*, *Tmem67^ΔCLE/+^* and *Tmem67^ΔCLE/ΔCLE^* lateral brain ventricles show highly elevated active Δ-catenin levels (indictive of elevated canonical Wnt signaling) in the *Tmem67* KO but not in the *Tmem67^ΔCLE/ΔCLE^*mouse brains which show a normal Wnt signaling signature. (**J**) Immunostaining for FOP (red) shows basal body apical surface localization is undisrupted and comparable in *Tmem67* KO and *Tmem67^ΔCLE/ΔCLE^* mouse brain ventricles. **** indicates a p-value < 0.0001, *** p-value < 0.001, ** p-value < 0.01, * p-value < 0.05 in Student’s unpaired t-test. Error bars indicate Mean ± S.D. in **B** and **E** and Mean ± S.E.M. in **G**. Scale bars in **A** are 10µm and 2µm and 10µm in **C**, **D, I** and **J**.

Since the *Tmem67^ΔCLE/ΔCLE^* mice phenocopied the *Tmem67*-null mice, we reasoned that further investigation of this model would yield new insights into the mechanism by which *TMEM67* mutations cause ciliopathies. The cysteine rich domain (CRD), present in the TMEM67 extracellular domain shed by the ADAMTS9-mediated cleavage, is highly homologous to the CRDs of the Wnt ligand binding Frizzled receptor [26], and TMEM67 has been implicated in the modulation of both canonical and non-canonical Wnt signaling pathways [24, 26–29, 55]. In particular, the TMEM67 extracellular domain is required for the activation of the non-canonical Wnt signaling pathway, forming a direct complex with Wnt5a and the non-canonical Wnt signaling receptor ROR2, and is required for phosphorylating ROR2 upon Wnt5a treatment [27, 28]. First, we tested whether the non-cleavable TMEM67- ΔCLE form could transduce non-canonical Wnt signaling by analyzing ROR2 phosphorylation *in vitro* (**Fig. 7H**). We found that, upon Wnt5a treatment, *Tmem67^ΔCLE/ΔCLE^* MEFs phosphorylated ROR2, similar to the control *Tmem67^ΔCLE/+^* MEFs, and reduced active β-catenin levels, a readout of the canonical Wnt signaling pathway. Second, to investigate the Wnt signaling signature *in vivo*, we analyzed the ependymal cells lining the brain ventricles of Wt, *Tmem67*-null, and *Tmem67^ΔCLE/ΔCLE^* mice. Active β-catenin staining revealed elevated active β-catenin staining in the *Tmem67*-null brains, presumably due to the loss of the non-canonical Wnt signaling pathway, but in contrast, normal levels of active β-catenin staining were seen in the *Tmem67^ΔCLE/ΔCLE^* brains (**Fig. 7I**). Immunostaining for the basal body marker FOP revealed unaltered apical localization of the basal bodies in *Tmem67^ΔCLE/ΔCLE^* and *Tmem67-null* brain ventricles (**Fig. 7J**).

Immunostaining for active β-catenin and F-actin upon Wnt3a or Wnt5a treatment of MEFs revealed increased active β-catenin staining and its nuclear localization upon activation of the canonical Wnt signaling pathway, and increased F-actin staining upon activation of the non-canonical Wnt signaling pathway, in both *Tmem67^ΔCLE/+^* and *Tmem67^ΔCLE/ΔCLE^* MEFs (**Fig. 8A-B**). qRT-PCR analysis of canonical and non-canonical Wnt signaling markers also revealed that *Tmem67^ΔCLE/ΔCLE^* MEFs transduced both signaling pathways as expected (**Fig. 8C-D**). To our surprise, we found that *Tmem67* transcription itself was regulated by canonical and non-canonical Wnt signaling pathways in Wt and *Tmem67^ΔCLE/ΔCLE^* MEFs (**Fig. 8E**).

**Figure 8:**
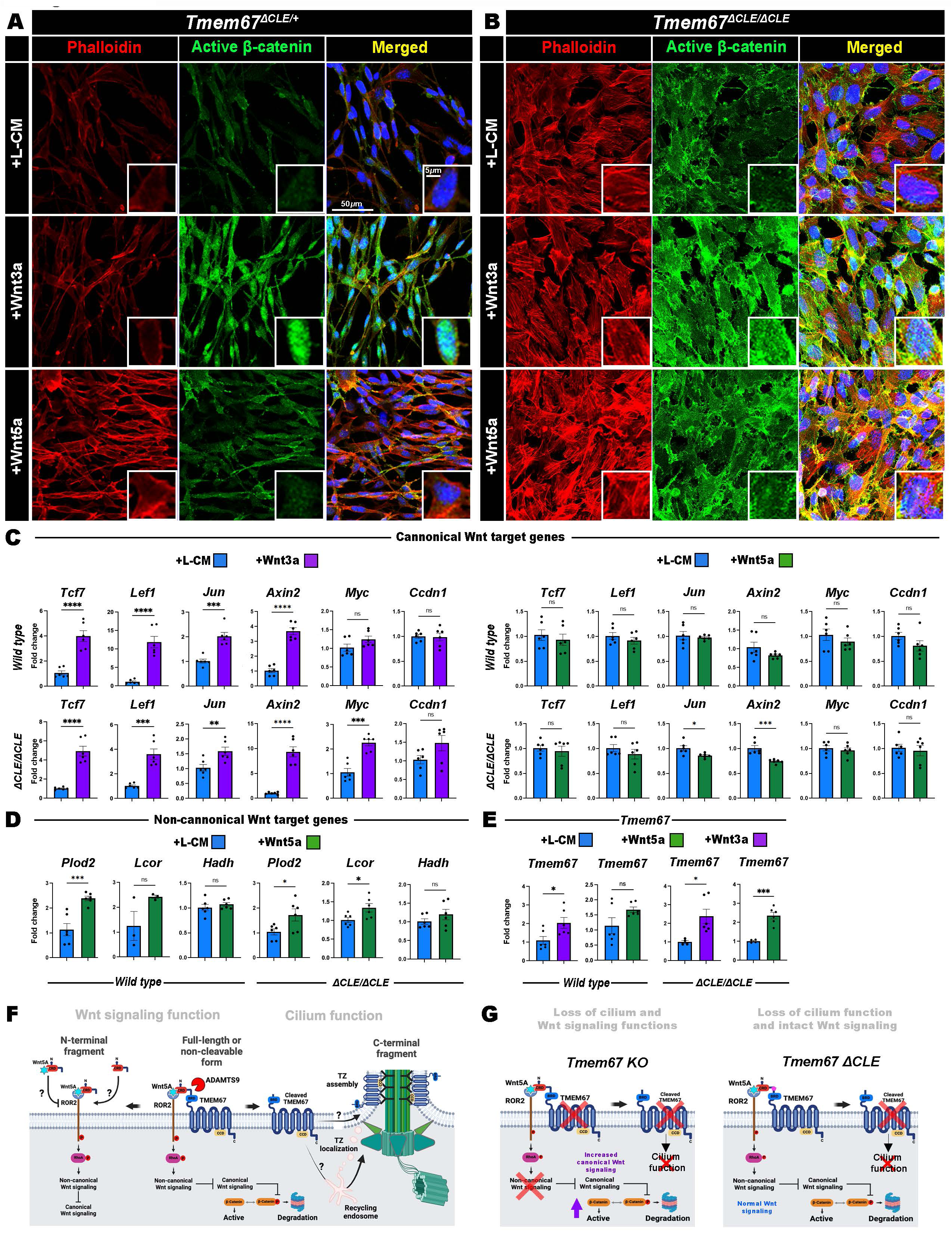
ADAMTS9-mediated TMEM67 cleavage regulates two distinct functions of TMEM67 in two distinct cellular compartments. (**A-B**) *Tmem67^ΔCLE/ΔCLE^* MEFs respond to canonical Wnt signaling stimulation (+Wnt3A) and show increased active β-catenin staining (green) and nuclear localization (inserts), while non-canonical Wnt signaling stimulation (+Wnt5a) causes increased F-actin staining (Phalloidin, red), similar to control *Tmem67^ΔCLE/+^* MEFs. (**D**) qRT-PCR analysis of *Wild type* and *Tmem67^ΔCLE/ΔCLE^* MEFs show canonical Wnt signaling stimulation (+Wnt3A, purple bars), upregulates the expression of known canonical Wnt target genes while non-canonical Wnt signaling stimulation (+Wnt5A, green bars) did not in comparison to L-cell conditioned medium treatment (blue bars). (**E**) qRT-PCR analysis of *Wild type* and *Tmem67^ΔCLE/ΔCLE^* MEFs treated with Wnt5A (green bars) show upregulation of some of the known non-canonical Wnt target genes. (**F**) qRT-PCR analysis of *Wild type* and *Tmem67^ΔCLE/ΔCLE^* MEFs treated with Wnt5A (green bars) or Wnt3A (purple bars) show *Tmem67* itself is regulated by both canonical and non-canonical Wnt signaling. (**G**) Cartoon depicting the proposed new model of how ADAMTS9-mediated TMEM67 cleavage may be regulating the abundance of two functional forms of TMEM67 which are active in two distinct cell membrane compartments. The N-terminally trimmed (TMEM67 Δ342) form localizes to the ciliary transition zone and regulates transition zone assembly while the full length form acts on regulating normal Wnt signaling on the cell surface. (**H**) Cartoon depicting the loss of both Wnt signaling and cilium functions in the *Tmem67* KO while *Tmem67^ΔCLE/ΔCLE^* mice transduce normal Wnt signaling but lose the ciliary TZ function. Scale bars in **A** and **B** are 100μm. **** indicates a p-value < 0.0001, *** p-value < 0.001, ** p-value < 0.01, * p-value < 0.05 in Student’s unpaired t-test. Error bars indicate Mean ± S.E.M. in **C-E**.

Combined, these results show that TMEM67 is indeed involved in regulating both normal Wnt signaling and ciliogenesis, supporting the findings of previous studies. Our novel findings provide the molecular mechanism by which TMEM67 performs these two functions (**Fig. 8F**). Cleavage of the protein by an extracellular matrix metalloproteinase generates the N-terminally cleaved TMEM67 (TMEM67 Δ342 form), which functions at the ciliary TZ and is required for normal assembly of the TZ necklace, while the full-length or the non-cleavable form of TMEM67 (TMEM67 ΔCLE) is required to regulate Wnt signaling. More importantly, these data also suggest that the molecular mechanisms of the ciliopathies caused by TMEM67 loss-of-function are due to loss of its TZ activity and not by its Wnt signaling activity, which could only be elucidated by the generation of the novel *Tmem67^1CLE/1CLE^* mice, which show defective ciliogenesis but normal Wnt signaling, whereas *Tmem67*-null mice are defective in both (**Fig. 8G**).

## DISCUSSION

Here we have uncovered an evolutionarily conserved cleavage motif present in the extracellular domain of the Meckel-Gruber syndrome protein TMEM67, revealing the molecular mechanism regulating the formation of two functional isoforms, by the extracellular metalloproteinase ADAMTS9. First, a C-terminal cleaved form, localized in the ciliary TZ, is involved in the formation of the TZ necklace and the MKS/B9 module assembly. We suggest that loss of this cleaved TMEM67 C-terminal fragment from the TZ, is the underlying disease-causing mechanism disrupted in ciliopathies, both in *Tmem67^ΔCLE/ΔCLE^* mice and, presumably, in humans with loss-of-function mutations. Second, the full-length non-cleaved form is essential for regulating normal Wnt signaling. ADAMTS9-mediated cleavage of TMEM67 regulates the abundance of each isoform and their respective functions and sub-cellular localizations (**Fig. 8F**). Whether the released N-terminal fragment containing the CRD can augment Wnt signaling acting long range and non-cell autonomously, remains to be determined in future studies. Our work suggests that the TMEM67 N-terminus needs to be proteolytically removed as a pre-requisite for TZ localization of the C-terminus and cilium assembly. This is equivalent to the “pro-domain” present in many secreted enzymes, which require proteolytic removal for their full functionality. Intracellular trafficking networks, signal peptides and post-translational modifications (PTMs) are all known to facilitate trafficking of molecules to the correct sub-cellular location. Our work has shown that removal of the TMEM67 N-terminus by ADAMTS9 is critical for the correct ciliary targeting of TMEM67, whereas the full-length protein exhibits a distinct localization pattern at the cell membrane and Golgi (**Fig. 1F**). Whether ADAMTS9-mediated cleavage causes TMEM67 C-terminal endocytosis and TZ localization *via* endocytic recycling vesicles emanating from the recycling endosome or causes lateral diffusion within the cell membrane during cilium assembly, remains unknown (**Fig. 8F**).

Previous studies from other groups have shown that proteolytic cleavage is a prerequisite for ciliary localization of the transmembrane proteins Polycystin-1 (PC1) and -2 (PC2), and mutations of the extracellular domain cleavage sites can also cause autosomal dominant polycystic kidney disease (ADPKD) in humans [56–60]. For PC1/PC2 complex to be effectively targeted to the cilium, PC1 is autoproteolytically cleaved at a G-protein coupled receptor cleavage site (GPS) during trafficking [61], but the exact molecules involved in the process remains unknown. Here we show that extracellular matrix metalloproteinases can facilitate transmembrane protein trafficking to the ciliary membrane. Since TMEM67 Δ342 only partially rescues ciliogenesis of *ADAMTS9*-null RPE1 cells, it is most likely that ADAMTS9 and its homologous sister protease ADAMTS20 may be involved in proteolytically cleaving other extracellular proteins essential for normal ciliogenesis. Since *Adamts9* and *Adamts20* are not ubiquitously expressed [62], whether additional extracellular proteases can cleave the TMEM67 extracellular domain in different tissues in the mammalian embryo also remains to be determined by future work.

Our comprehensive immunostaining analysis of 14 *bona fide* TZ proteins revealed 6 MKS/B9 proteins (TCTN1, TCTN2, TCTN3, TMEM237, CC2D2A and B9D2) were reduced or lost at the TZ in both *TMEM67* and *ADAMTS9* KO cells, while components of the NPHP module were mostly unaffected. Analysis of *Tmem67^ΔCLE/ΔCLE^* primary cilia and motile cilia by TEM revealed the TZ necklace is completely missing upon loss of TMEM67 cleavage and its localization to the TZ. Intriguingly, loss of TCTN1 from photoreceptor cilia also prevents ciliary necklace formation and the TZ gate-keeping activity in mice [63], highlighting the significant role played by the extracellular and transmembrane components of the transition zone. TZ modules occupy distinct positions along the axial plane of the TZ, and in previous characterization by super-resolution microscopy revealed that NPHP proteins exhibit a narrower diameter located towards the axonemal microtubules, while the MKS complex proteins occupy a wider region towards the ciliary membrane [64, 65]. Here we show the wider MKS/B9 proteins are lost or reduced in *TMEM67* KO cells, while TZ proteins belonging to the NPHP module are largely unaffected. We therefore propose first, that TZ assembly may be initiated in an inside out manner from the axonemal microtubules to the membrane, as proteins lost in *TMEM67* KO cells also exhibit a wider axial diameter. If the TZ was assembled from the membrane inwards, we would instead expect the loss of narrower, internal TZ proteins following loss of TMEM67. Second, TZ assembly may also be occurring in a modular fashion, where NPHP and MKS modules are assembled independently. Future in-depth studies probing this specific question are required to confirm this, however, and this study provides a basis to better understand the molecular hierarchy of ciliary TZ proteins in relation to TMEM67. Furthermore, the functional domains and motifs also relate to their location within the TZ. NPHP proteins, for example, are enriched in microtubule-binding domains and coiled-coil motifs in comparison to MKS proteins that comprise the cell membrane-binding (B9, C2), transmembrane, and extracellular domains. CEP290 and RPGRIP1L are enriched in coiled-coil motifs and are known to be coordinators of TZ assembly, with CEP290 interacting with proteins from both the NPHP and MKS modules [8, 42, 49, 66]. Our findings support the notion that TMEM67 plays a central role in anchoring the extracellular (TCTN1) and transmembrane components of the TZ (which forms the necklace), to the Y linkers, which remain unaffected in both *TMEM67* KO and *ADAMTS9* KO TZs. We speculate that the TMEM67 coiled-coil domain may act as a crucial molecular scaffold for its “outside-in” anchoring role at the TZ. A future study investigating a mutated CCD in TMEM67 Δ342 would provide valuable insight in answering this intriguing question.

In addition to ciliary TZ assembly, here we have demonstrated that *Tmem67^ΔCLE/ΔCLE^* MEFs maintain their ability to phosphorylate ROR2, indicating that non-cleaved full-length TMEM67 maintains functionality in the non-canonical Wnt pathway. The *in vivo* data, in comparison to *Tmem67*-null brains, confirms that TMEM67 indeed performs two discrete functions and is involved in Wnt signaling regulation as well. The generation of the *Tmem67^ΔCLE/ΔCLE^* mouse model as part of this study provided a unique opportunity to investigate this dual functionality of TMEM67 reported in previous work by many others, which has been an unresolved and highly debated question in the field. The findings from this study reveal that TMEM67 is not a *single functional protein* acting in two distinct cellular compartments and in two crucial cellular pathways simultaneously as previously thought, but in fact are *two functional isoforms* of a single gene product, acting in two distinct cellular compartments, and should be studied as such in the future.

Whether the released N-terminal CRD fragment (N-331) retains its Wnt signaling activity is unknown. We hypothesize that the TMEM67 N-331 may retain its bioactivity and may function similar to a soluble frizzled related protein (SFRP) [67], and it may have the ability to augment both canonical and non-canonical Wnt signaling activity as a recombinant molecule, acting long range and non-cell autonomously. Future work probing this question will answer this aspect of TMEM67 cleavage uncovered by this work.

TMEM67 is the most commonly mutated gene in MKS, but is also causative of NPHP, JBTS, RHYNS, and COACH syndromes. The dual functionality of TMEM67, driven by proteolytic cleavage, may explain the huge range and severity of ciliopathies linked to TMEM67 mutations ranging from fetal lethality to adults with mild liver fibrosis only. The N-terminal TMEM67 cleavage product identified here (TMEM67 N-331) and its potential use to augment dysregulated Wnt signaling in polycystic kidney disease and many other human diseases where altered Wnt signaling plays a central role [68–72], is an exciting new paradigm and byproduct uncovered by this study, which we plan on exploring further in future studies.

## ACKNOWLEDGMENTS

This work was funded by the National Institute of Health grant 5R01DK126804 and The Worcester Foundation grant awards to SN. This work utilized SEM, TEM and Ultramicrotomy equipment that was purchased from the NIH awards S10RR021043, S10OD025113, and SI0OD021580, awarded to the UMass Chan Medical School EM core facility. A TissueGnostics SL slide scanner used in the study was funded by a Massachusetts Life Science Center, Bits to Bytes grant award. SB acknowledges support from a Wellcome Trust Clinical Training Fellowship (4Ward North Clinical PhD Academy, 203914/Z/16/Z). The research was supported by MRC project grants (MR/M000532/1 and MR/T017503/1) to CAJ, and a bilateral BBSRC-SFI project grant (BB/P007791/1) to CAJ and OB. We thank Professors George Witman and Piali Sengupta and Dr. Yuqing Hou for their valuable input and discussions on the work, Dr. Greg Hendricks and Keith Redding of the UMass Chan Medical School EM core facility for their outstanding electron microscopy expertise, and Drs. Christina Baer and Jill McConnell of the SCOPE imaging center for their help in confocal and super-resolution microscopy.

## AUTHOR CONTRIBUTIONS

M.A., S.F, K.L., K.R., O.B. and S.N. conceived and designed the experiments. M.A., S.N., S.F, K.L., S.B., K.R., and M.S. performed the experiments. S.B. and C.A.J. provided reagents. M.A., K.L., and S.N. wrote the manuscript. All authors read, edited and approved the manuscript.

## COMPETING INTERESTS

None of the authors have any conflicts to declare.

---

**Supplemental Figure 1:**
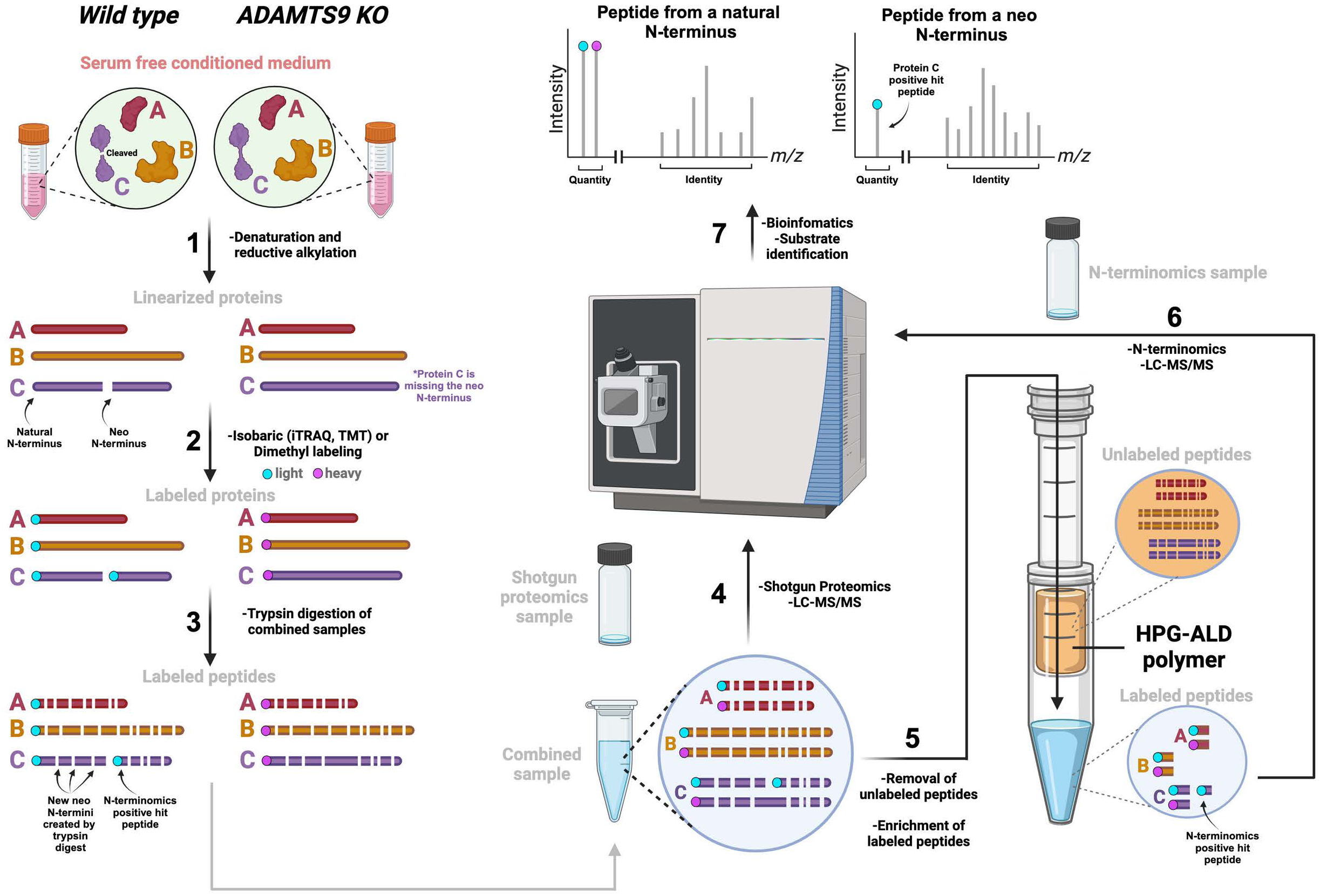
Summary of the TAILS N-terminomics technique used to identify TMEM67 as a novel substrate of ADAMTS9. Protein C (purple) is shown as an example substrate of ADAMTS9 and is cleaved only in the Wt sample generating a neo N-terminus, which is absent in the *ADAMTS9* KO sample. The turquoise or magenta labels represent two forms of labeling either utilizing isobaric tags or heavy/light dimethyl labeling techniques.

**Supplemental Figure 2:**
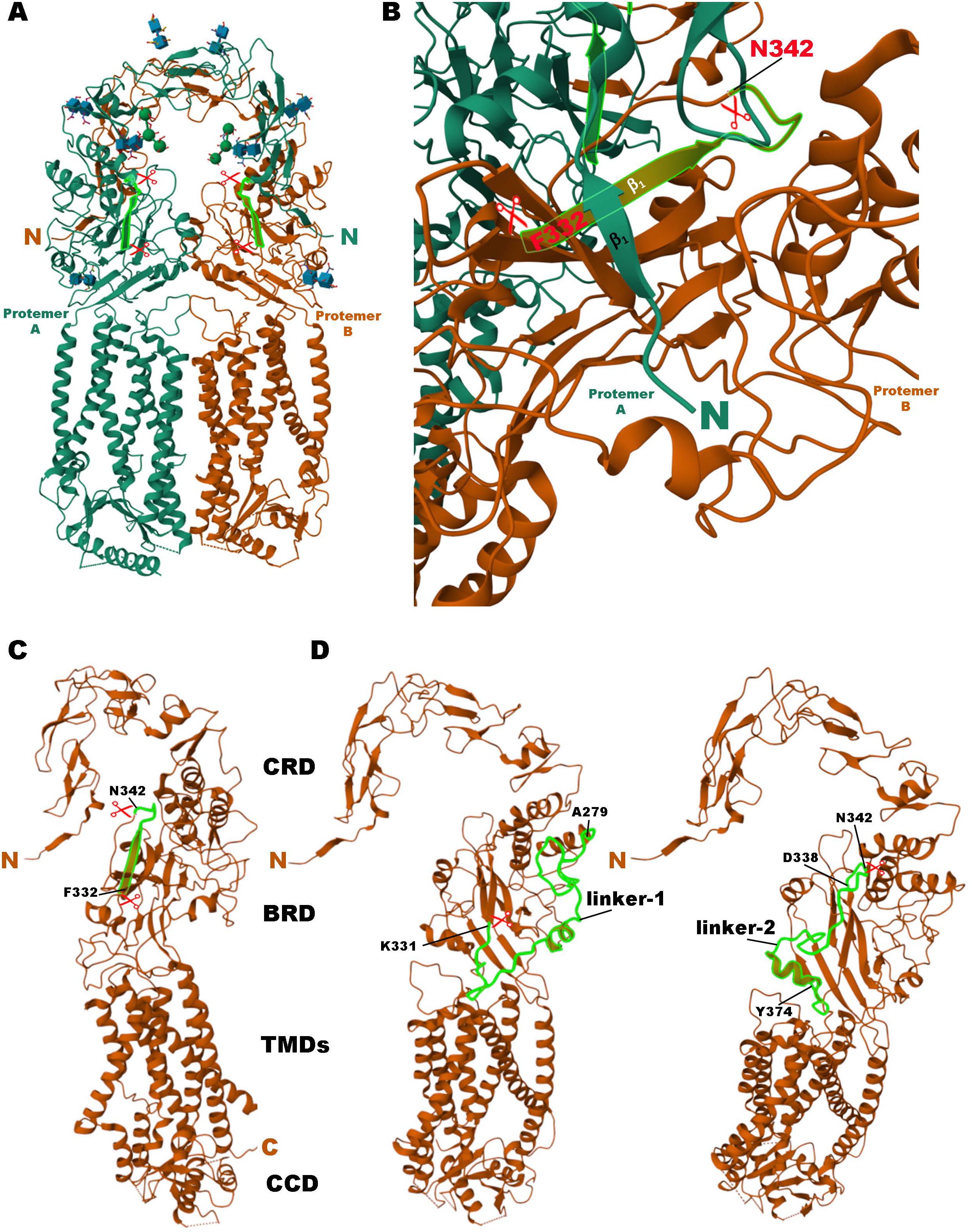
Mapping of the ADAMTS9 cleavage sites and the linkers 1 and 2 in the TMEM67 cryo-EM resolved structure. (**A**) TMEM67 dimer structure showing the two ADAMTS9 cleavage sites in each protomer (red scissors) and the 11 amino acid peptide released by the cleavage (green highlight) is shown. ADAMTS9-mediated cleavage of a dimerized TMEM67 form would still release a CRD-CRD dimer as the cleavage occurs at the point of the first interface of the two protomers as shown. (**B**) Higher magnification view of the Protomer A (green) N-terminal end coding for the β1 loop of its CRD interfacing with the Protomer B (orange) β1 loop of its BRD is shown. The two ADAMTS9 cleavage residues in Protomer B is shown by the red scissors. The 11 amino acid Protomer B sequence released by the cleavage is highlighted in luminescent green. Concurrent cleavage occurring in Protomer A releases the dimerized CRD. (**C**) Cryo-EM structure of a single TMEM67 protomer is shown marking the cleavage residues (red scissors) and the 11 amino acid peptide fragment (green). (**D**) Cryo-EM structure of a single TMEM67 protomer marking the 52 amino acid linker-1 (A^279^ to K^331^) containing cleavage site 1 and the 36 amino acid linker-2 (D^338^ to Y^374^) containing the second cleavage site are shown. Each linker sequence is highlighted in green. CRD, cysteine rich domain; BRD, β-sheet rich domain; TMDs, transmembrane domains; CCD, coiled-coil domain.

**Supplemental Figure 3:**
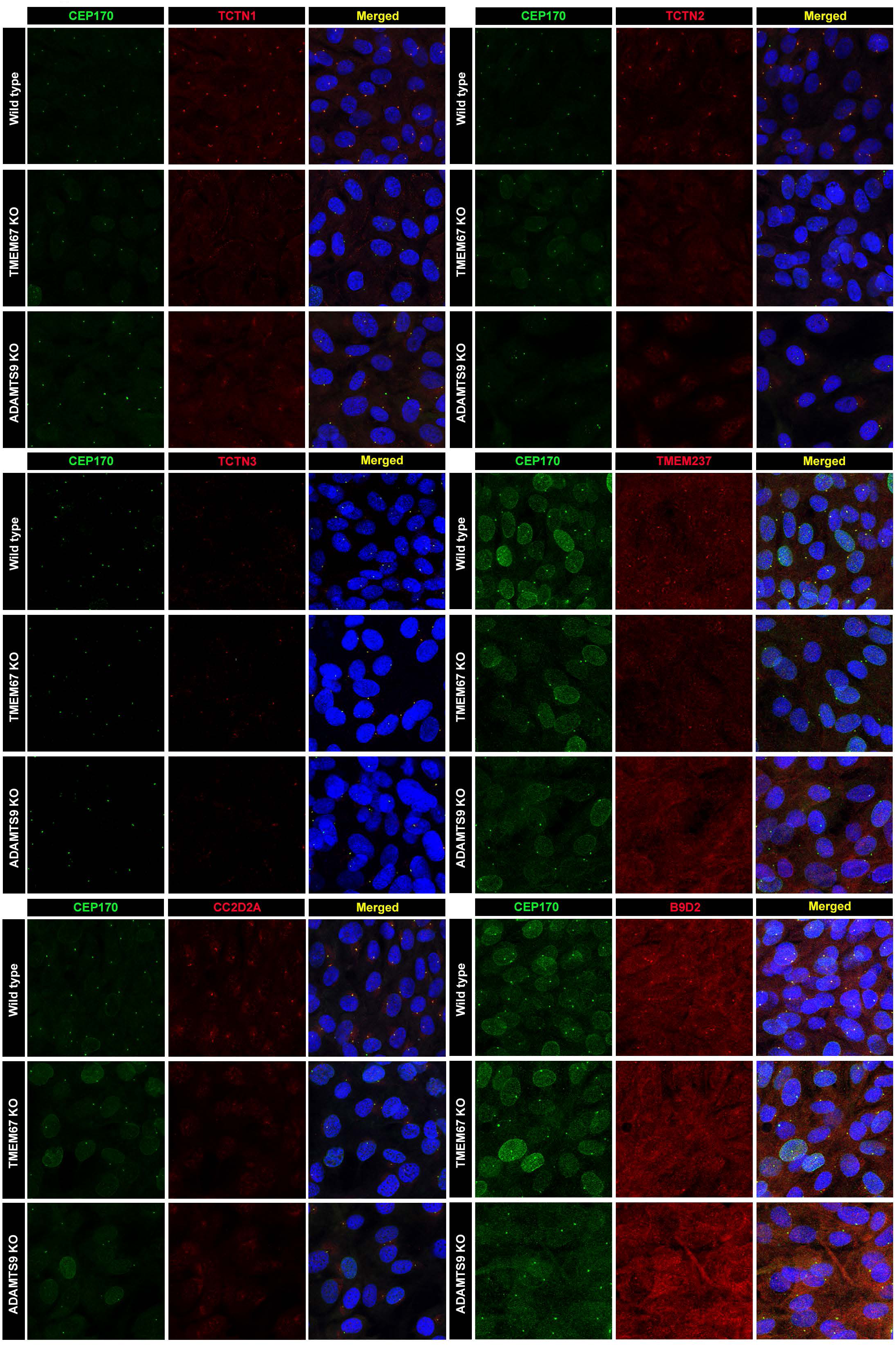
Significantly affected transition zone molecules in both *TMEM67* KO and *ADAMTS9* KO RPE-1 cells. Conventional high resolution confocal microscopy images labeling the indicated transition zone molecule (red) and the mature basal body marker (CEP170) in serum starved RPE-1 cells are shown. Scale bar is 100μm.

**Supplemental Figure 4:**
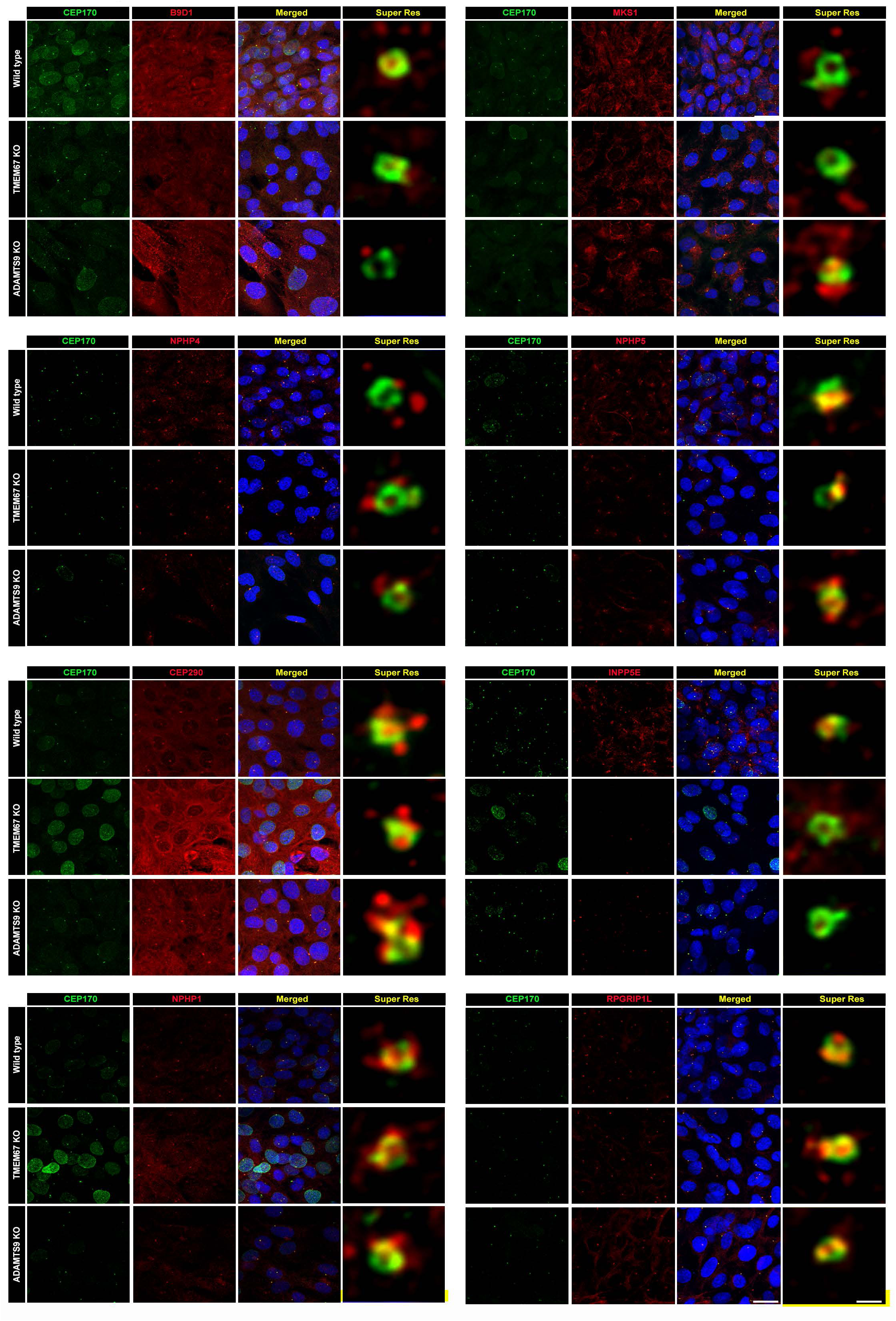
Transition zone molecules significantly unaffected in both *TMEM67* KO and *ADAMTS9* KO RPE-1 cells. Conventional and super resolution confocal microscopy images labeling the indicated transition zone molecule (red) and the mature basal body marker (CEP170) in serum starved RPE-1 cells are shown. Scale bars are 100μm and 500nm.

**Supplemental Figure 5:**
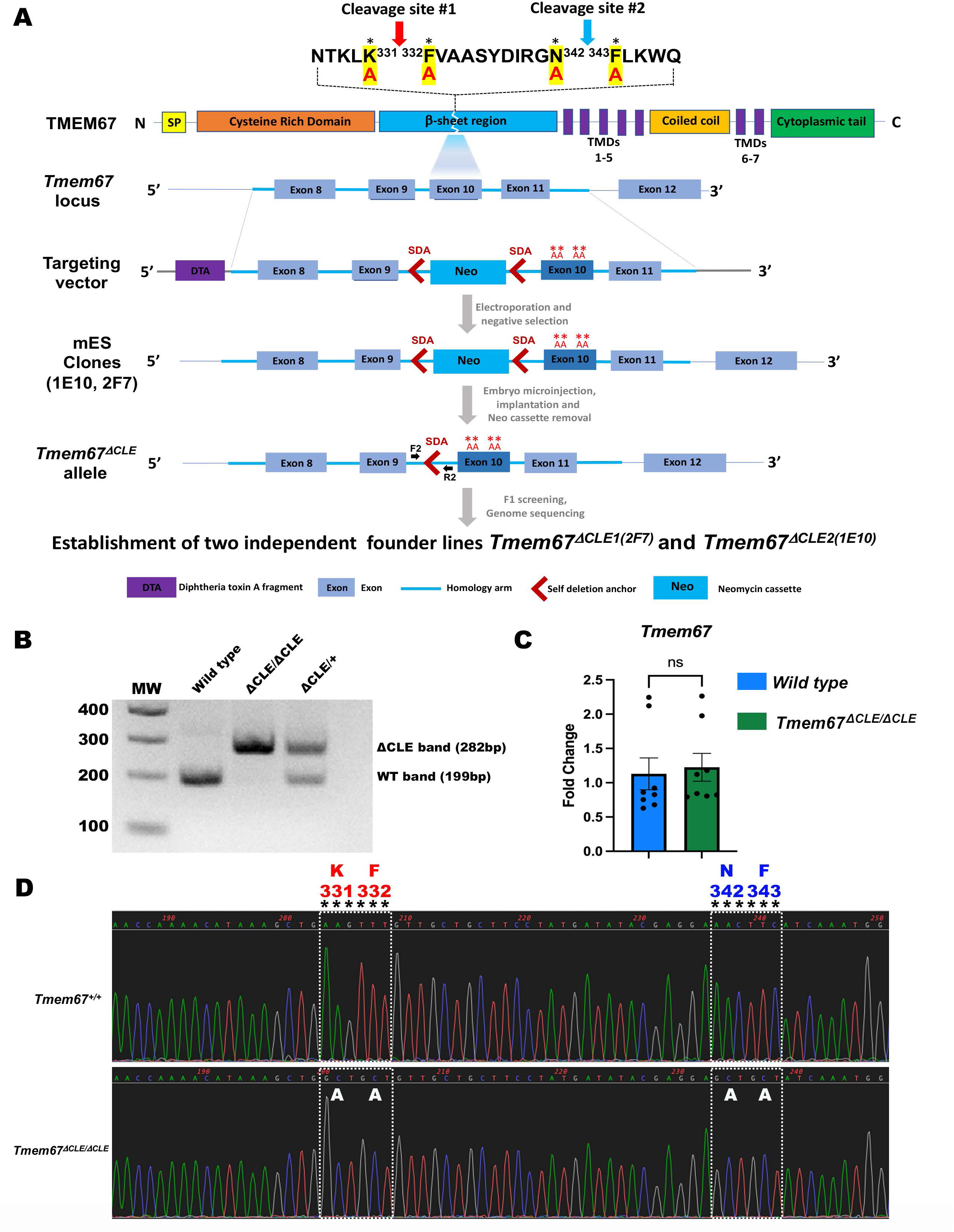
Generation of the novel non-cleavable *Tmem67^ΔCLE/ΔCLE^* mouse model. (**A**) Strategy used for generating two independent founder mouse lines *Tmem67^ΔCLE1^* and *Tmem67^ΔCLE2^* from two genetically engineered mouse embryonic stem cell clones (2F7 and 1E10), and the location of the genotyping primers are shown. (**B**) A DNA gel showing Wt and *ΔCLE* bands generated from the PCR amplification of genomic DNA using primers F2 and R2 from each genotype indicated is shown. (**C**) qRT-PCR for *Tmem67* transcript levels show no significant difference in Wt and *Tmem67^ΔCLE/ΔCLE^* kidneys. (**D**) Sanger sequencing results of the *Tmem67* exon 10 shows both cleavage site residues have been successfully mutated into Alanine in *Tmem67^ΔCLE/ΔCLE^* mice.

**Supplemental Figure 6:**
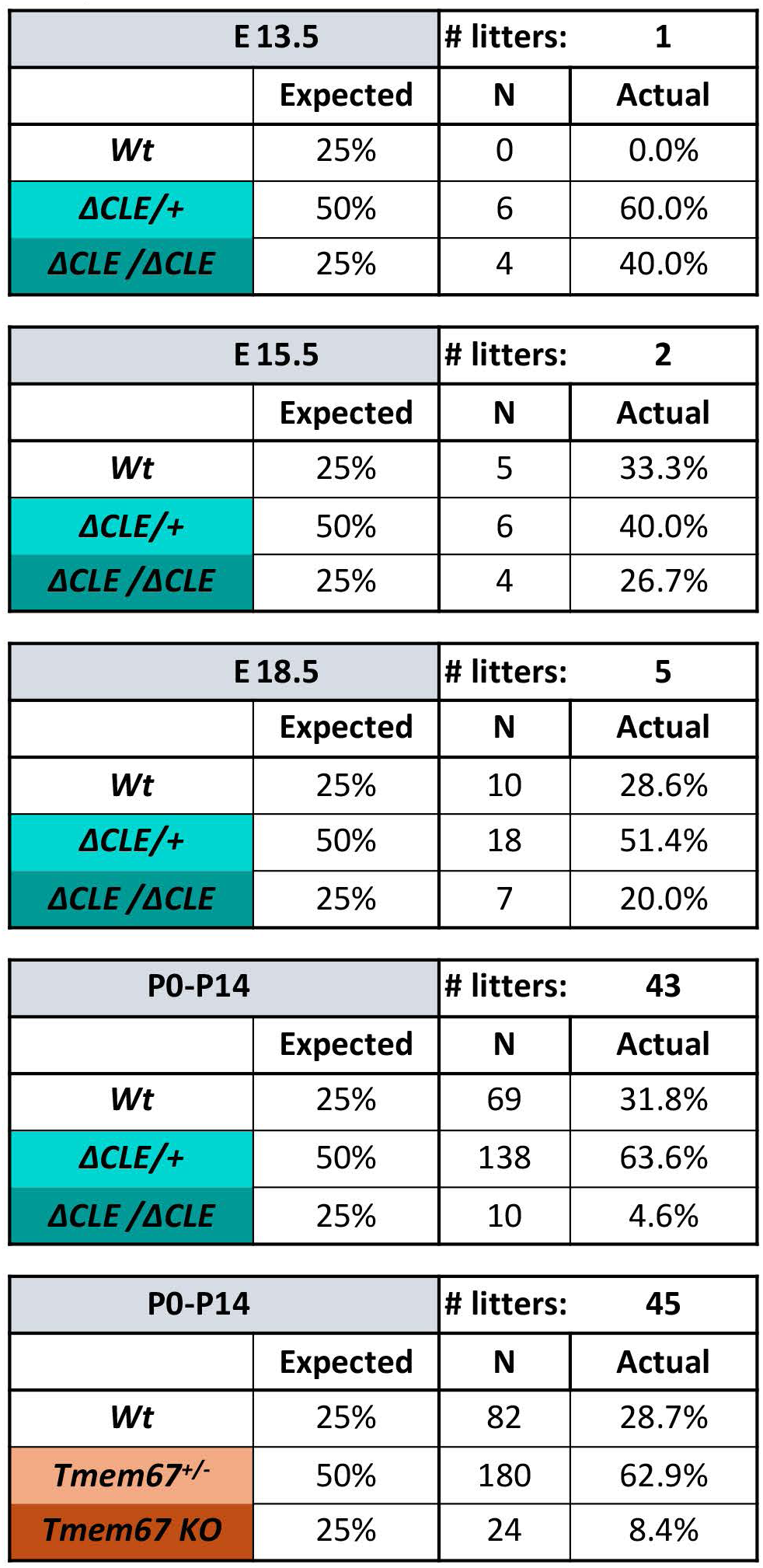
Expected and observed mouse numbers of *Tmem67^ΔCLE/+^* and *Tmem67^+/-^* mouse breeding.

**Supplemental Figure 7:**
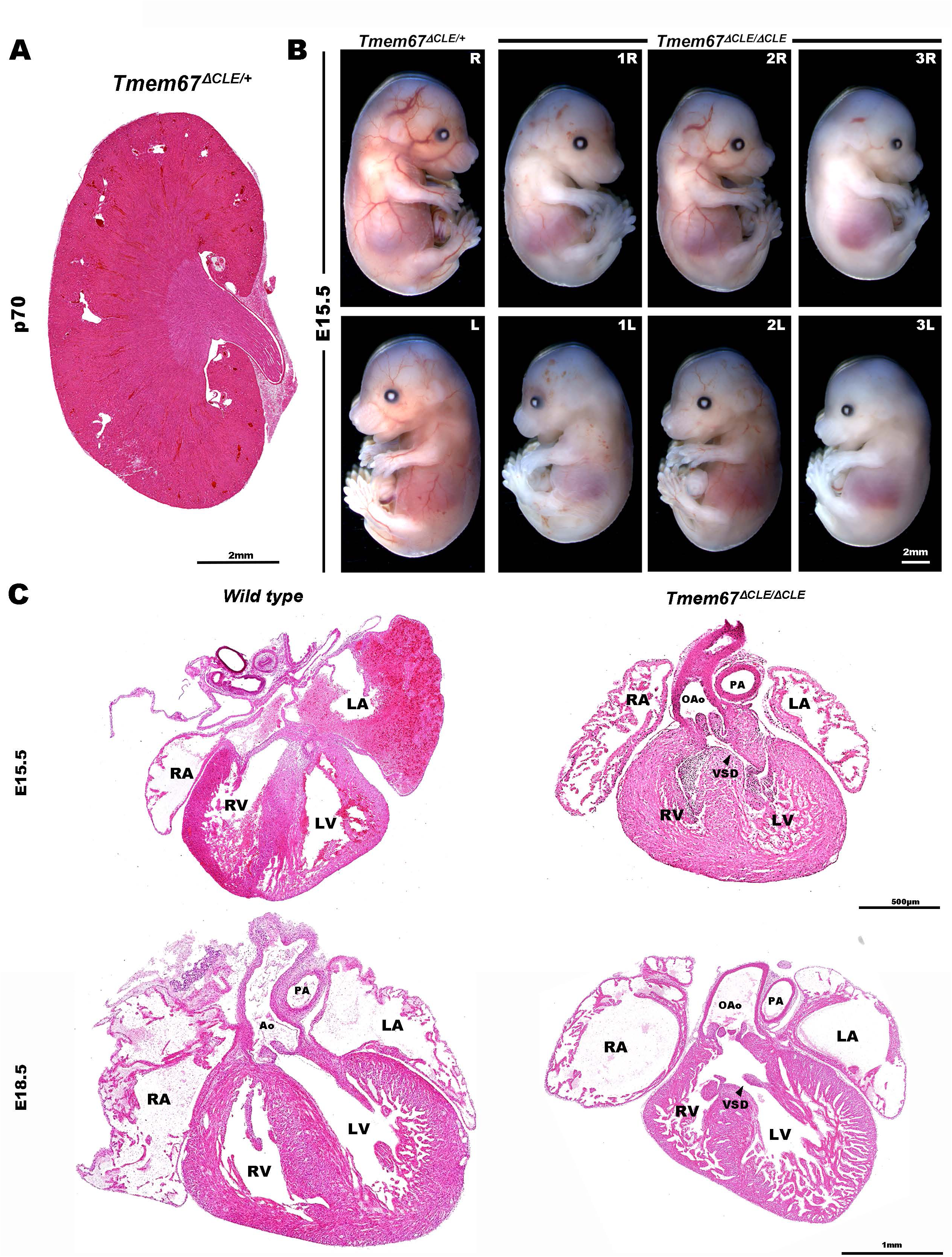
Cardiac and vascular phenotypes in *Tmem67^ΔCLE/ΔCLE^*mouse embryos. (**A**) Hematoxylin and Eosin stained kidney section from a 10 week old (p70) *Tmem67^ΔCLE1/+^* mouse is shown. Heterozygous (ΔCLE/+) mice are asymptomatic and have normal life spans and do not develop cystic kidneys. (**B**) Embryos harvested on day 16 (E15.5) show hemorrhaging and loss of vascular circulation in many *Tmem67^ΔCLE/ΔCLE^* embryos, leading to embryonic lethality. Left and right sides of three homozygous embryos and one heterozygous embryo are shown. (**C**) Hematoxylin and Eosin stained E15.5 and E18.5 heart sections show an overriding aorta, ventricular-septal defect and defective myocardial compaction in *Tmem67^ΔCLE/ΔCLE^*hearts. RA, right atrium; LA, left atrium; RV, right ventricle; LV, left ventricle; Ao, aorta; OAo, overriding aorta; PA, pulmonary artery; VSD, ventricular septal defect. Scale bars in **A** and **B** are 2mm, 500μm and 1mm in **C**.

**Supplemental Figure 8:**
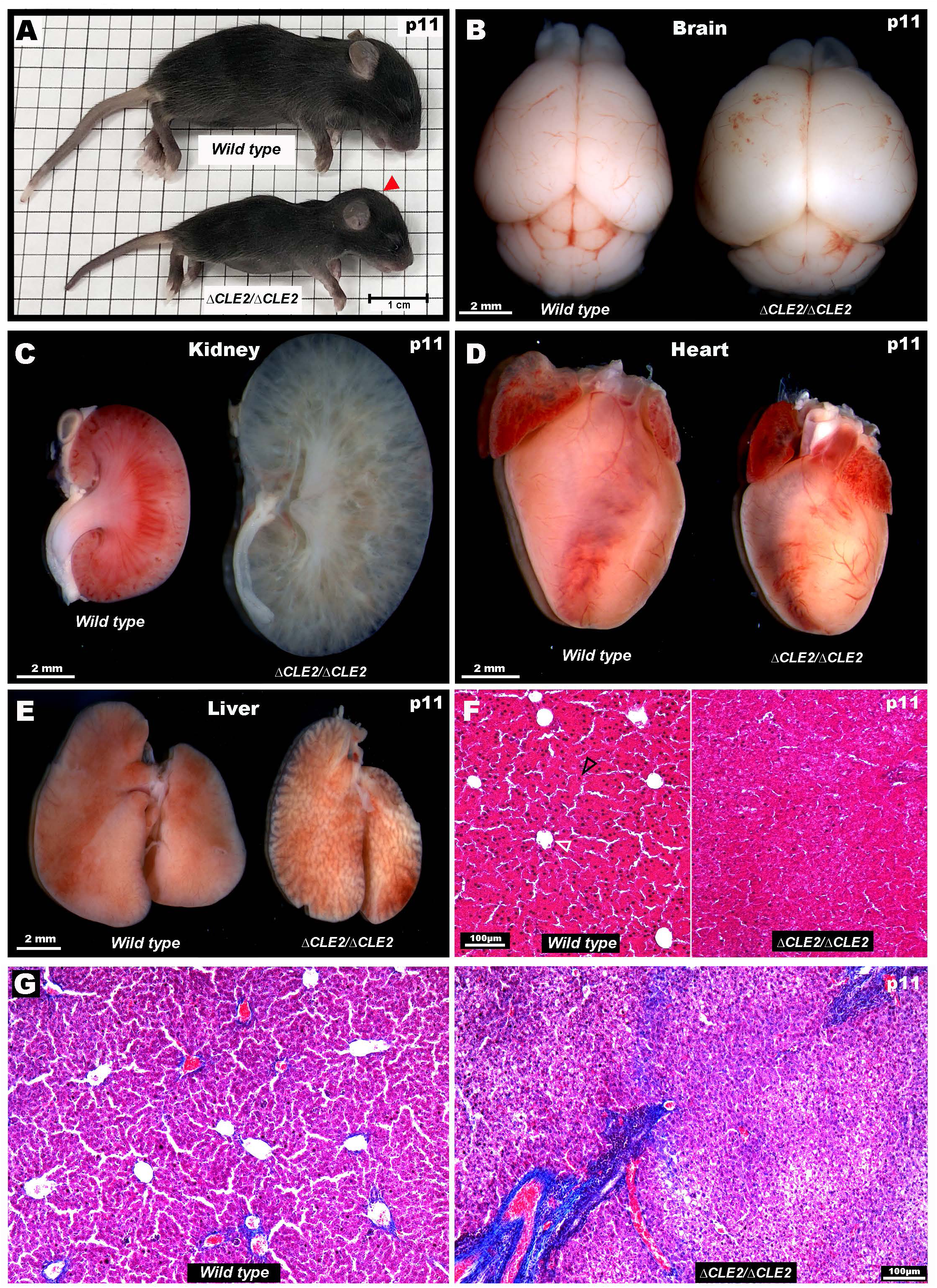
Homozygous mice for *Tmem67^ΔCLE1^* and *Tmem67^ΔCLE2^*are phenotypically identical to one another and phenocopy *Tmem67*^-/-^ mice causing ciliopathies in multiple organs. (**A**) A male Wt mouse and a same sex littermate *Tmem67^ΔCLE2/ΔCLE2^* mouse at postnatal day 11 (p11) is shown. *Tmem67^ΔCLE2/ΔCLE2^* mice fail to thrive postnatally and are smaller than their Wt and heterozygote littermates. A dome-shaped head (red arrowhead), indicative of hydrocephaly, can be seen in the homozygous mutants as early as postnatal day 9. (**B**) Dissected mouse brains from p11 Wt and *Tmem67^ΔCLE2/ΔCLE2^* littermate mice are shown. Mutant brains are wider and shorter compared to Wt brains due to develop hydrocephaly formation. (**C**) Bisected kidneys from Wt and *Tmem67^ΔCLE2/ΔCLE2^* mice at p11, photographed post PFA fixation are shown. *Tmem67^ΔCLE2/ΔCLE2^* mice develop very large polycystic kidneys and do not survive past p14. (**D**) *Tmem67^ΔCLE2/ΔCLE2^* mice also show defective heart rotation and smaller hearts compared to Wt littermates. Ventral view of the dissected hearts post fixation are shown. (**E**) Dissected whole livers from *Tmem67^ΔCLE2/ΔCLE2^* mice are morphologically distinguishable from Wt littermates and are relatively smaller. (**F**) Hematoxylin and Eosin staining of liver sections show hepatic portal vein branching morphogenesis and hepatocyte (black arrowhead) differentiation is severely impaired in the mutant livers. White arrowhead indicate central veins. (**G**) Masson’s trichrome staining of liver sections show onset of fibrous expansion and deposition of lipid droplets in *Tmem67^ΔCLE2/ΔCLE2^* mice. Scale bar in **A** is 1cm, 2mm in **B-E** and 100μm in **F- G**.

**Supplemental Table-1:**
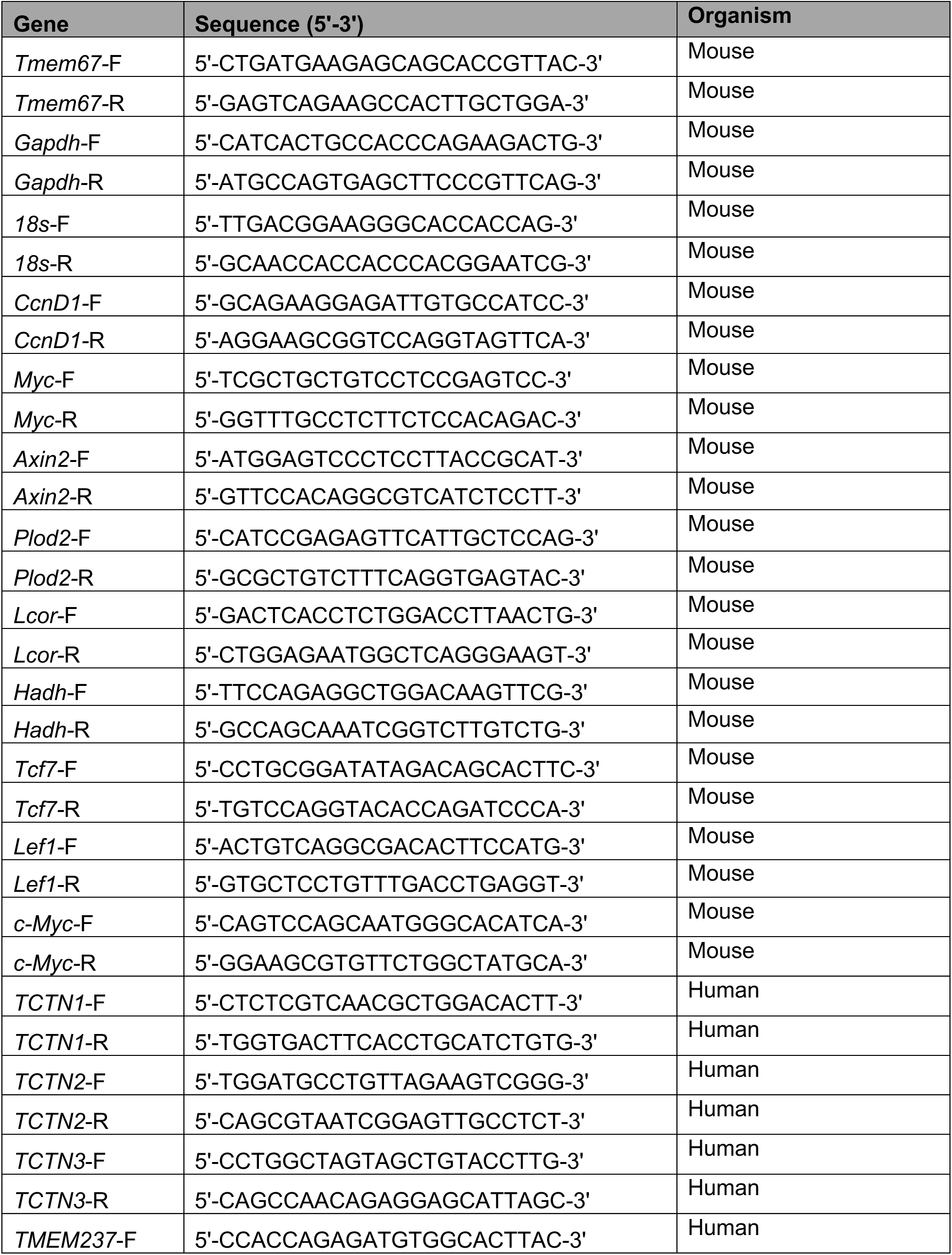
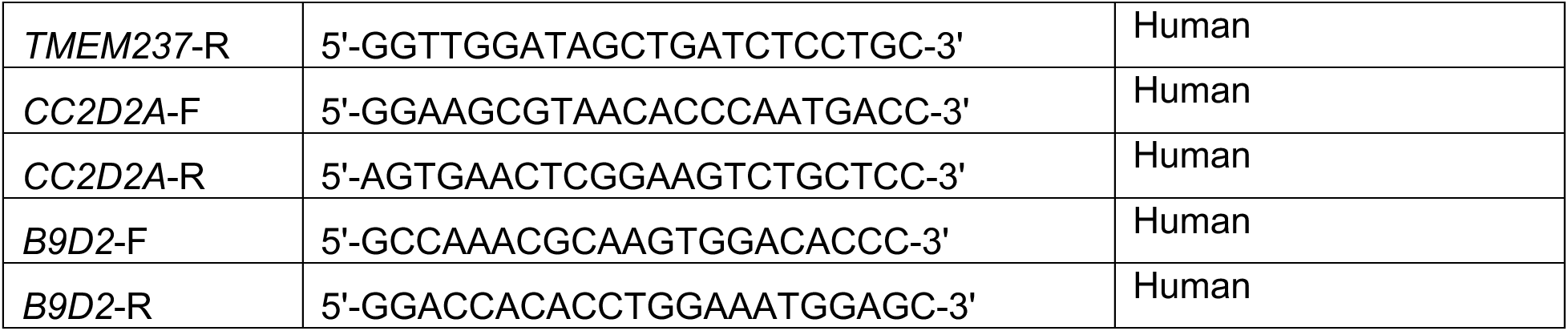
Mouse and Human qRT-PCR primers used in this study.

**Supplemental Table-2:**
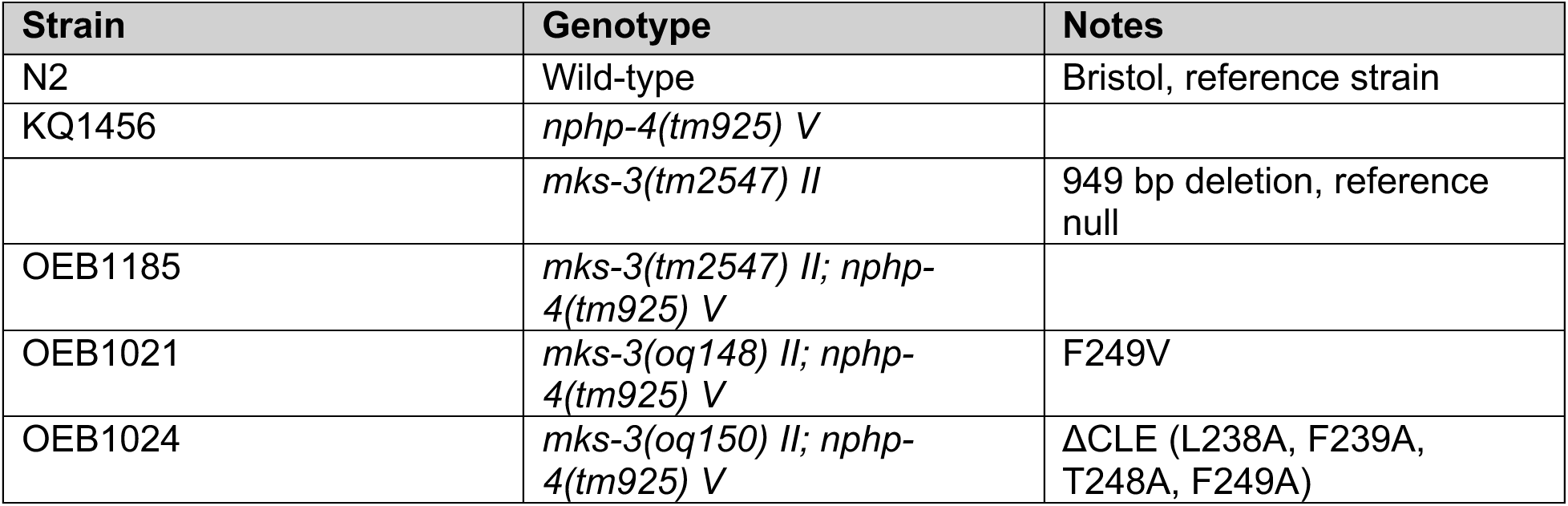
*C. elegans* strains used in this study.

| Strain | Genotype | Notes |
| --- | --- | --- |
| N2 | Wild-type | Bristol, reference strain |
| KQ1456 | *nphp-4(tm925) V* |  |
|  | *mks-3(tm2547) II* | 949 bp deletion, reference null |
| OEB1185 | *mks-3(tm2547) II; nphp-4(tm925) V* |  |
| OEB1021 | *mks-3(oq148) II; nphp-4(tm925) V* | F249V |
| OEB1024 | *mks-3(oq150) II; nphp-4(tm925) V* | ΔCLE (L238A, F239A, T248A, F249A) |

**Supplemental Table-3:**
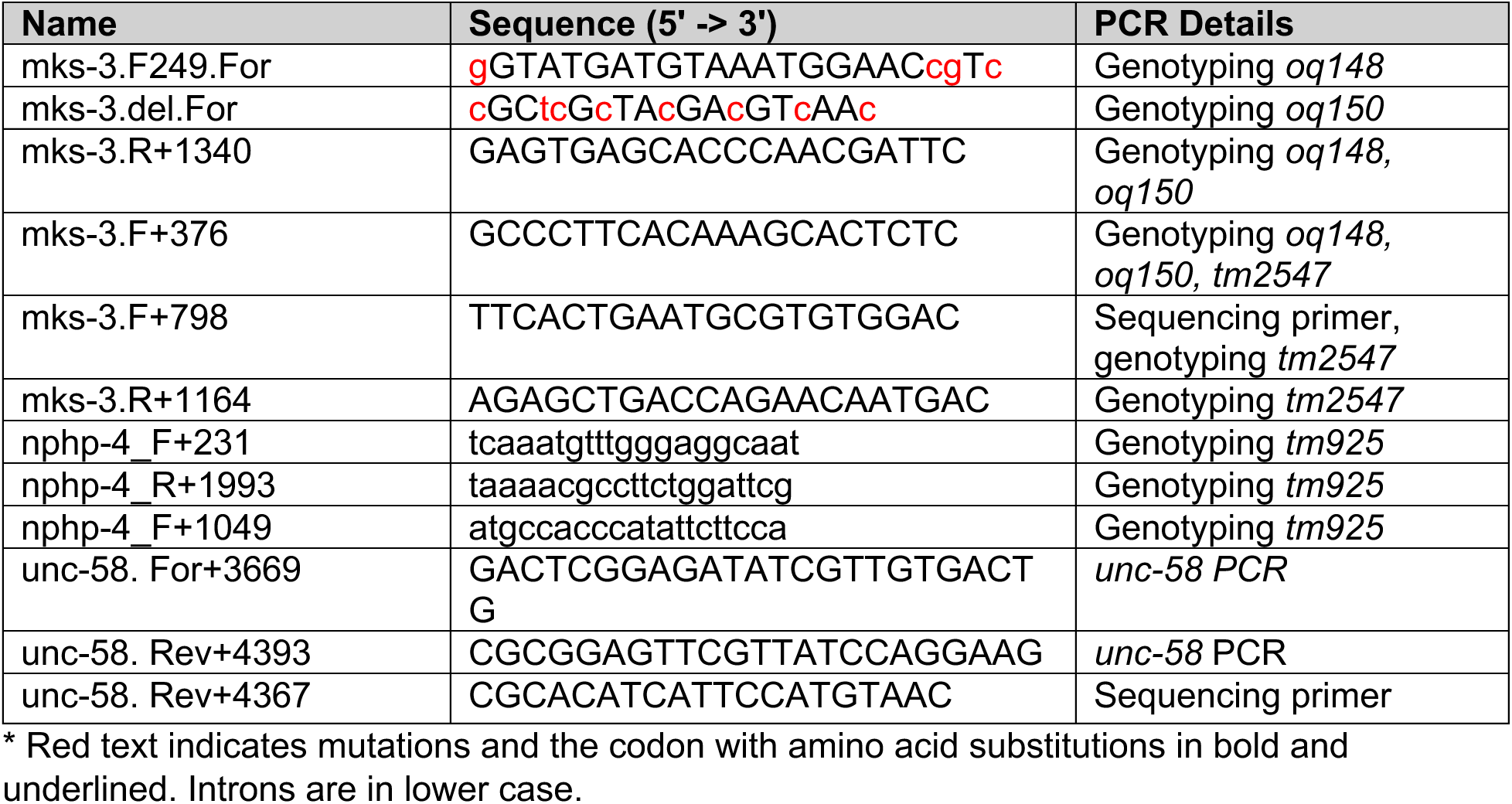
*C. elegans* primers used in this study.

**Supplemental Table-4:**
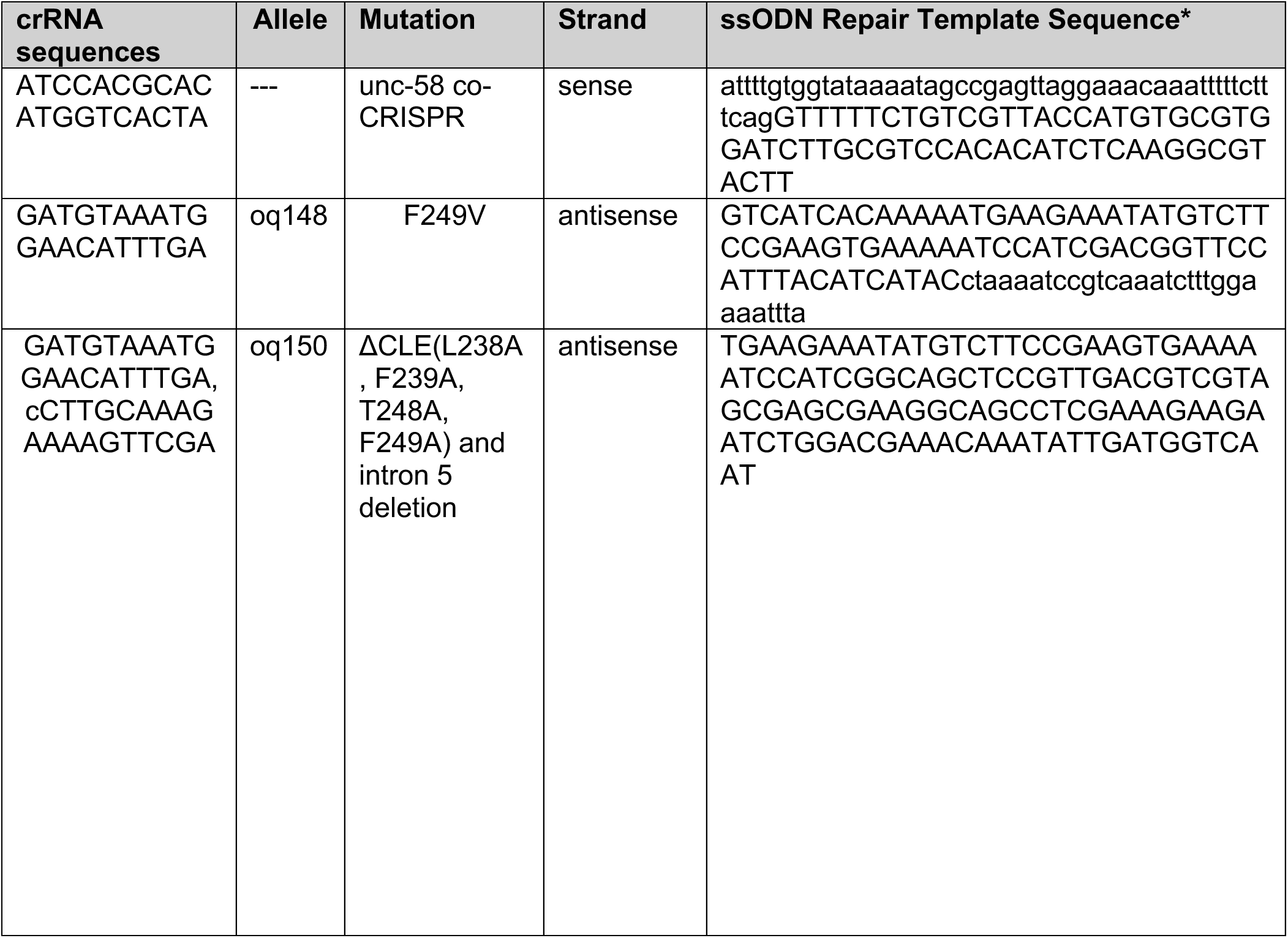
*C. elegans* crRNA and repair template sequences used in this study.

## KEY RESOURCE TABLE

| Resource | Source | Identifier (RRID) |
| --- | --- | --- |
| **Antibodies** |  |  |
| Goat anti-ROR2 (WB, 1:1000) | R&D Systems | AB_2180118 |
| Mouse anti-ROR2 (WB, 1:1000) | Proteintech | AB_2918662 |
| Mouse anti-GAPDH (WB, 1:1000) | Millipore | AB_2107426 |
| Rabbit active β-catenin (WB, 1:1000; IF, 1:200) | Cell Signaling Technology | AB_11127203 |
| Alexa Fluor 568 Phalloidin (IF, 1:400) | ThermoFisher Scientific | A12380 |
| Rabbit anti-TMEM67 (MKS3) (IF, 1:500) | Proteintech | AB_10638441 |
| Mouse anti-CEP170 (IF, 1:400) | ThermoFisher Scientific | AB_2533502 |
| Mouse anti-γ tubulin (IF, 1:200) | Abcam | AB_297920 |
| Rabbit anti-acetylated α-tubulin (IF, 1:500) | Cell Signaling Technology | AB_10544694 |
| Rabbit anti-TCTN1 (WB, 1:1000, IF, 1:400) | Proteintech | AB_10644442 |
| Rabbit anti-TCTN2 (WB, 1:1000, IF, 1:400) | Proteintech | AB_2202406 |
| Rabbit anti-TCTN3 (WB, 1:1000, IF, 1:400) | Proteintech | AB_10638442 |
| Rabbit anti-TMEM237 (WB, 1:1000, IF, 1:400) | ThermoFisher Scientific | AB_2648665 |
| Rabbit anti-CC2D2A (WB, 1:1000, IF, 1:400) | Proteintech | AB_2879063 |
| Rabbit anti-B9D1 (IF, 1:400) | ThermoFisher Scientific | AB_2808685 |
| Rabbit anti-B9D2 (WB, 1:1000, IF, 1:400) | Proteintech | AB_2878981 |
| Rabbit anti-MKS1 (IF, 1:400) | Proteintech | AB_10637856 |
| Rabbit anti-CEP290 (IF, 1:400) | ThermoFisher Scientific | AB_2780038 |
| Rabbit anti-RPGRIP1L (IF, 1:400) | Proteintech | AB_2923607 |
| Rabbit anti-NPHP1 (IF, 1:400) | Thermo Fisher Scientific | AB_795024 |
| Rabbit anti-NPHP4 (IF, 1:400) | Proteintech | AB_10640302 |
| Rabbit anti-NPHP5 (IQCB1) (IF, 1:400) | Atlas Antibodies | AB_2677806 |
| Rabbit anti-INPP5E (IF, 1:400) | Proteintech | AB_2167120 |
| Rabbit anti-FGFT1OP (FOP) (IF, 1:400) | Proteintech | AB_2103362 |
| Goat anti-Mouse Alexa Fluor 488 (IF, 1:500) | ThermoFisher Scientific | AB_2536161 |
| Goat anti-Rabbit Alexa Fluor 488 (IF, 1: 500) | ThermoFisher Scientific | AB_2633280 |
| Goat anti-Mouse Alexa Fluor 568 (IF, 1: 500) | ThermoFisher Scientific | AB_144696 |
| Goat anti-Rabbit Alexa Fluor 568 (IF, 1: 500) | ThermoFisher Scientific | AB_143157 |
| Donkey anti-Mouse, VRDye 549 (WB, 1:10,000) | Li-COR Biosciences | 926-54012 |
| Goat anti-Mouse, VRDye 680 (WB, 1:10,000) | Li-COR Biosciences | AB_2651128 |
| Donkey anti-Rabbit, VRDye 549 (WB, 1:10,000) | Li-COR Biosciences | 926-54013 |
| Goat anti-Rabbit, VRDye 800 (WB, 1:10,000) | Li-COR Biosciences | AB_2651127 |
| **Bacterial and Viral strains** |  |  |
| DH5α | NEB | C2987H |
| Simian virus 40 large T antigen (SV40LT) | Greg Pazour lab | N/A |
| **Chemicals, reagents and media** |  |  |
| DMEM/F-12 | Gibco | 10565018 |
| Penicillin/Streptomycin | Gibco | 15140-122 |
| Hygromycin B | Invitrogen | 10687010 |
| FBS | Gibco | A52567-01 |
| Opti-MEM | Gibco | 31985-070 |
| PEI MAX | Polysciences | 24765 |
| 16% Formaldehyde | Electron Microscopy Sciences | 15710 |
| Trypsin-EDTA | Gibco | 25300-054 |
| Trizol RNA extraction reagent | Invitrogen | 15596026 |
| Bullseye Real Time qPCR master mix EvaGreen | MidSci | BEQPCR-S |
| Intercept (TBS) Blocking Buffer<br>Normal Goat Serum | LiCor<br>Invitrogen | 927-60001<br>31872 |
| **Experimental models: Cell lines** |  |  |
| hTERT-RPE1<br>TMEM67 KO RPE-1<br>ADAMTS9 KO RPE-1<br>L-Cells<br>L-Wnt3A<br>L-Wnt5A | ATCC<br>Colin Johnson lab<br>Sumeda Nandadasa lab<br>ATCC<br>ATCC<br>ATCC | CVCL_4388<br>N/A<br>N/A<br>CVCL_4536<br>CVCL_0635<br>CVCL_0636 |
| **Experimental models: Mouse lines** |  |  |
| *Tmem67*<sup>ΔCLE/ΔCLE</sup><br>*Tmem67*<sup>+/-</sup> (C57BL/6NJ-*Tmem67*<sup>em1(IMPC)J</sup>/Mmjax)<br>C57BL6/J *Wild type* | This study<br>Jackson Laboratories<br>Jackson Laboratories | N/A<br>MMRRC_051248-JAX<br>IMSR_JAX:000664 |
| **Experimental models: Worm lines** |  |  |
| N2 *Wild type* worms | Caenorhabditis Genetics Center |  |
| *nphp-4(tm925) V* | Ashrafi lab |  |
| *mks-3(tm2547) II* | Blacque lab |  |
| *mks-3(tm2547) II; nphp-4(tm925) V* | Blacque lab |  |
| *mks-3(oq148) II; nphp-4(tm925) V* (F249V) | This study | N/A |
| *mks-3(oq150) II; nphp-4(tm925) V* [ΔCLE (L238A, F239A, T248A, F249A)] | This study | N/A |
| **Recombinant DNA** |  |  |
| TMEM67 Full-length pCDNA3.1 Myc/His | Colin Johnson lab | N/A |
| TMEM67 S1 pCDNA3.1 Myc/His | Sumeda Nandadasa lab | N/A |
| TMEM67 S2 pCDNA3.1 Myc/His | Sumeda Nandadasa lab | N/A |
| TMEM67 S1+S2 pCDNA3.1 Myc/His | Sumeda Nandadasa lab | N/A |
| TMEM67 N-331 pSecTag2C Myc/His | Sumeda Nandadasa lab | N/A |
| TMEM67 Δ342 pSecTag2C Myc/His | Sumeda Nandadasa lab | N/A |
| TMEM67 F343V pCDNA3.1 Myc/His | Sumeda Nandadasa lab | N/A |
| TMEM67 K329T pCDNA3.1 Myc/His | Sumeda Nandadasa lab | N/A |
| TMEM67 L349S pCDNA3.1 Myc/His | Sumeda Nandadasa lab | N/A |
| **Software and Algorithms** |  |  |
| Image J FIJI<br>LI-COR Acquisition (Ver 1.0)<br>Leica Application Suite X (LAS-X)<br>GraphPad Prism (Ver 10.3.0)<br>Bio-Rad CFX Maestro (Ver 2.3) | NIH<br>Li-COR Biosciences<br>Leica Microsystems<br>GraphPad<br>Bio-Rad | SCR_003070<br>N/A<br>SCR_013673<br>SCR_002798<br>N/A |

## Notes

### Competing Interest Statement

The authors have declared no competing interest.

